# Food for the Empire: dietary pattern of Imperial Rome inhabitants

**DOI:** 10.1101/2020.01.23.911370

**Authors:** Flavio De Angelis, Sara Varano, Andrea Battistini, Stefania Di Giannantonio, Paola Ricci, Carmine Lubritto, Giulia Facchin, Luca Brancazi, Riccardo Santangeli-Valenzani, Paola Catalano, Valentina Gazzaniga, Olga Rickards, Cristina Martínez-Labarga

**Affiliations:** Centre of Molecular Anthropology for Ancient DNA Studies, University of Rome Tor Vergata. Via della Ricerca Scientifica 1, 00133 Rome, Italy; Collaborator Servizio di Antropologia, Soprintendenza Speciale Archeologia, Belle Arti e Paesaggio di Roma, Rome, Italy; Dipartimento di Scienze e Tecnologie Ambientali, Biologiche e Farmaceutiche Università degli Studi della Campania “Luigi Vanvitelli”, Via Vivaldi 43, 81100 Caserta, Italy; Dipartimento di Studi Umanistici, Università degli Studi Roma Tre, Via Ostiense 234-236, 00146 Rome, Italy; Scuola di Dottorato in Archeologia, Dipartimento di Scienze dell’Antichità, Sapienza Università di Roma, Piazzale Aldo Moro 5, 00185 Rome, Italy; former Servizio di Antropologia, Soprintendenza Speciale Archeologia, Belle Arti e Paesaggio di Roma, Rome, Italy; Unità di Storia della Medicina e Bioetica, Sapienza University of Rome, Viale dell’Università 34, 00185 Rome, Italy

**Keywords:** Imperial Rome, Diet, Carbon and nitrogen stable isotopes

## Abstract

This paper aims to provide a broad diet reconstruction for people buried in archaeologically defined contexts in Rome (1^st^-3^rd^ centuries CE), in order to combine archaeological and biological evidence focusing on dietary preferences in Imperial Rome.

A sample of 214 human bones recovered from 6 funerary contexts were selected for carbon and nitrogen stable isotope analysis. The baseline for the terrestrial protein component of the diet was set using 17 coeval faunal remains recovered from excavations at Rome supplemented by previously published data for the same geographic and chronological frames.

δ^13^C ranges from −19.95‰ to −14.78‰, whereas δ^15^N values are between 7.17‰ and 10.00‰. The values are consistent with an overall diet mainly based on terrestrial resources. All the human samples rely on a higher trophic level than the primary consumer faunal samples.

Certainly, C_3_ plants played a pivotal role in the dietary habits. However, C_4_ plants also seem to have been consumed, albeit they were not as widespread and were not always used for human consumption. The environment played a critical role also for Romans of lower social classes. The topographical location determined the preferential consumption of food that people could obtain from their neighborhood.

## INTRODUCTION

Imperial Rome was one of the largest cities of Europe (Scheidel, 2007) and feeding its population was a serious concern for political authorities. Demographic surveys witness a peak in both urban and suburban Roman populations during the Imperial Age (1^st^-3^rd^ centuries CE, herein indicated by the capitalized word “Empire,” whereas the uncapitalized word “empire” refers to the geographical boundaries, as suggested by Boatwright (2012)), revealing that about one million people lived in the city or within 50 km. Nearly 17% of the Italian population was concentrated in just 5% of Italian territory (Morley, 1996; Scheidel, 2001) and this affected public health as well as administrative and social organization (Dyson, 2010).

Roman authorities began to step in the food supply of the city in the mid-Republican period. The introduction of grain distribution by *C. Sempronius Gracchus* in 123 BCE is considered the first legal provision for supplying the citizens of Rome. According to this rule, each legal resident was entitled to receive a monthly allotment of basic foods at a discounted price or even for free. Because wheat supplied most of the calories citizens consumed, the government focused its interventions in the wheat market, especially for the poor, although meat and oil were also distributed in later years. Eligibility for the food allotment required an ever-watchful eye by the authorities. From the second half of the first century BCE, the names of those entitled to receive the *frumentatio* were recorded in dedicated registers. However, eligibility for the provision could also be acquired by donation or by the purchase of the *frumentaria* card, the tablet on which the name of the eligible citizen was engraved.

In the Principate, the *Annona* (the grain supply) was a critical element of the relationship between the Emperor and the citizens and this office was headed by a powerful political leader. Beyond the imperial estates’ production, the empire collected tax grain primarily in Sicily and Africa. The obtained stock was distributed at the *frumentationes*, which fed a large part of the population but not its entirety. The basic conditions for accessing the public supply were Roman citizenship, residence in Rome, being male, and being of legal age, though there were many exceptions (Johnson et al., 2013). Of course, the food requirements of Rome could not be fulfilled only by the central distribution of supplementary grain and Roman social stratification in the city and suburbs created many related problems.

Archaeological evidence suggests the area outside the city walls, the *Suburbium*, was inhabited both by poor people, who could not afford the city lifestyle, as well as by the upper strata of Roman society, who wanted to spend their lives outside the unhealthy urbanized environment (Champlin, 1982). This liminal area between the city and the open countryside also included marginal industries excluded from the city for religious or public safety reasons, such as landfills, quarry pits, brickmaking facilities, and funerary areas (Killgrove and Tykot, 2013, Catalano, 2015). Movements between the *Urbs* and the *Suburbium* were frequent and, according to Witcher, the permanent Rome-ward migration from the countryside helped to maintain the population size of Rome (Scheidel, 2007), that was granted by people with different origins and cultural features (Killgrove and Tykot, 2013, Antonio et al., 2019), including their dietary habits.

Roman diet was and continues to represent a fertile area of investigation, and the historic and iconographical record provides a great deal of evidence of the variety of foodstuffs available to at least some of the Roman populace. Food was a popular motif in the decoration of Roman estates, where wealthy Romans enjoyed a fully catered lifestyle, especially in rooms associated with food consumption, such as kitchens and dining rooms (Yardley, 1991). However, these luxury items were undoubtedly mainly produced by and for the upper social stratum, representing less than 2% of the population.

There is still little evidence about the diet of commoners living in the empire, especially in the area of Rome (Killgrove and Tykot, 2013). The broadest discussion of the diet of ancient Romans is provided by primary sources, such as novels and artworks (Purcell, 2003; Wilkins and Hill, 2006).

Despite primary information about diet provided by texts describing foods and ancient recipes (Garnsey, 1999), no direct evidence could be clearly identified concerning the food consumption of the common people and poor people of Rome. Grain would have been the base of their diet and carbohydrates from grains would have accounted for about 70% of their daily energy intake (Delgado et al., 2017). Grain was used in a variety of recipes, mainly as bread or *puls*, a grain pottage that could also be mixed with vegetables, meat, and cheese (Garnsey, 1999). Accordingly, cereals were widely cultivated in the empire, and consistent importation came from areas like Sicily and Egypt. The commercial value of grain was determined by the Edict of Diocletian, which set the maximum price of wheat, barley, and millet. Remarkably, the role of millet is still not completely understood, and it might have been mainly used for livestock fodder rather than for human sustenance (Spurr, 1983). The pivotal role of cereals in the Empire is also attested to by evidence concerning Roman skill in ensuring a continuous supply of those foodstuffs through diverse agricultural practices, artificial farming techniques, and food preservation methods (De Ligt, 2006). Along with the cereal backbone, a wide variety of vegetables, fruits, and legumes were eaten (or drunk, as in the case of wine) by Romans (Garnsey,1999; Prowse, 2001).

Certainly, meat represented a critical element of an individual’s food consumption: livestock breeding and trade were extremely widespread in the Roman world (Kron, 2002; MacKinnon, 2004) and the main sources of meat were goats, sheep, lambs, and pigs (Brothwell and Brothwell, 1998; MacKinnon, 2004). Furthermore, the role of fish in the Empire is unclear as this foodstuff was alternatively seen as an expensive or a common food (Purcell, 2003) in various ecological contexts. According to Galen, marine fish were more highly valued than freshwater fish, and their consumption in ancient Rome increased with *garum*, the staple fish sauce.

Information about the Roman diet could also be provided by mounting archaeobotanical evidence found at roughly coeval sites, such as the floral remains from Pompeii and Herculaneum (Rowan, 2017). Similarly, recovered faunal remains suggest the types of meat and fish available to Romans (King, 1999; Cool, 2006; Prowse et al., 2004; 2005).

The evaluation of human bone remains recovered in archaeological contexts could provide an even clearer glimpse into the lives of the people who lived and died in Rome. Indeed, human bones play a critical role in the evaluation of a community’s subsistence strategy through carbon and nitrogen stable isotope analysis of bone collagen (De Niro, 1985; Ambrose and Norr, 1993; O’Brien, 2015). Thus, the spread of investigations into ancient diets using carbon and nitrogen stable isotope analysis in recent years has begun to reveal dietary habits in several regions of the Rome area (O’Connel et al., 2019; Killgrove and Tykot 2017; Killgrove and Montgomery 2016; Killgrove and Tykot 2013; Bruun 2010; Crowe et al., 2010; Killgrove and Tykot, 2013; Rutgers et al., 2009; Prowse et al., 2004; Prowse 2001), though the assessment of a significant sample of commoners who were buried (and perhaps lived) in the nearby *Suburbium* is still far from being proficiently accounted for.

Dietary information represents a critical source of knowledge into complex societies such as ancient Rome as it has now been established that customs around food are a key tool for understanding the relationship between humans and their cultural and natural environment in the past (Smith, 2006). Therefore, this paper aims to provide a broad diet reconstruction for people buried in archaeologically defined contexts in Rome, in order to combine archaeological and biological evidence as well as recent excavation results focusing on dietary preferences in Imperial Rome.

## MATERIALS AND METHODS

### SAMPLE

A sample of 214 human bones (Table 1) recovered from 6 funerary contexts (Figure 1) were selected for carbon and nitrogen stable isotope analysis. The visible preservation status of the skeletons was the leading inclusion criterion for the recruitment. Information on sex and age at death for each individual were available from previous studies (Catalano, 2015), in which the results of osteometric, and paleopathological analyses were reported.

**Figure 1:**
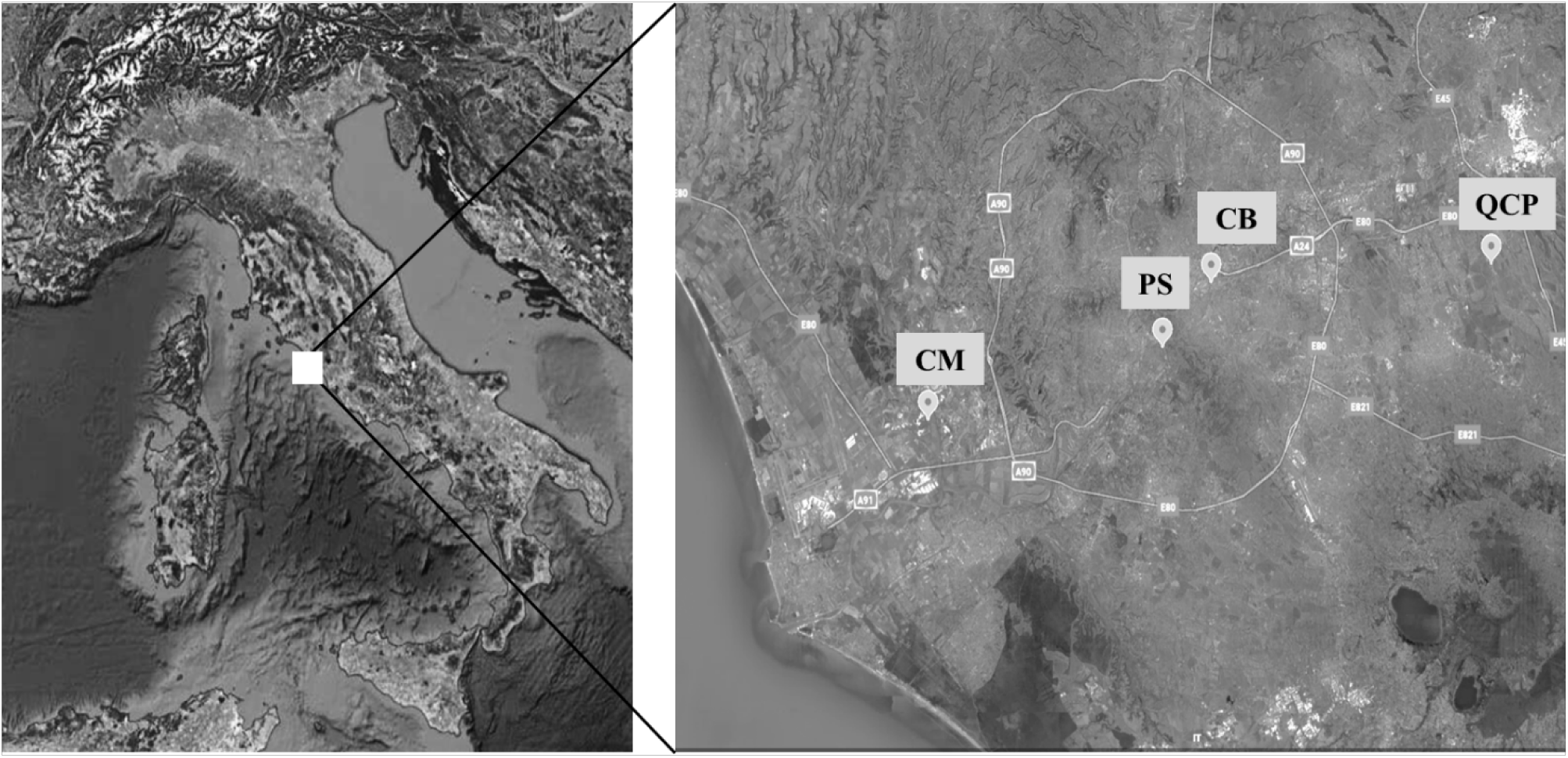
Topographical locations of the funerary areas. CM: Castel Malnome, PS: Via Padre Semeria; CB: Casal Bertone; QCP: Quarto Cappello del Prete.

**Table 1:** Sample size for each funerary context.

| **Funerary context** | **Code** | **Sample size** |
| --- | --- | --- |
| **Castel Malnome** | CM | 79 |
| **Casal Bertone Mausoleum** | CBM | 26 |
| **Casal Bertone Area Q** | CBQ | 20 |
| **Casal Bertone Necropolis** | CBN | 19 |
| **Via Padre Semeria** | PS | 30 |
| **Quarto Cappello del Prete** | QCP | 40 |

The necropolis of Castel Malnome was excavated in the southwestern suburbs (Catalano et al 2010; Catalano et al 2013). The sex ratio and juvenile index value, along with osteological suggestions related to musculoskeletal stress markers, push to consider that the funerary area was related to the salt flats unearthed close to the necropolis, where its living community might have worked (Caldarini et al., 2015).

The burial ground of Casal Bertone was set in the eastern suburbs close to the Aurelian walls, in proximity to a large productive area related to an ancient tannery (*fullonica*) (Musco et al, 2008). The funerary context was archaeologically subdivided into three sections: a mausoleum, a necropolis, and an area, named Area Q, contiguous to the productive area. The demographic profile of the mausoleum and necropolis communities allows us to consider them as a unique population (De Angelis et al., 2015), and the analysis of skeletal stress markers suggests the population from both areas could have been engaged in work at the *fullonica*. Conversely, the demographic profile of Area Q is significantly dissimilar to the others and is characterized by a peculiar distribution of mortality, in which 48% were in the 0-6 years age range. This has been explained by the hazardous environmental conditions in Area Q, evidenced by the presence of pathological alterations likely caused by infectious diseases (De Angelis et al., 2015).

Quarto Cappello del Prete necropolis was established in the extreme eastern suburbs of Rome, along the Via Prenestina, near the ancient city of Gabii (Musco et al, 2010). Monumental structures, such as a circular basin and a nymphaeum, were found at the site, and the graves were located along the edges of a pool and in a hypogeum. More than 70% of the buried people were infants and juveniles; 50% of them were in the 0-6 years age range, and more than a half of them seem to have suffered from dysmorphic alterations (De Angelis et al., 2015).

The funerary area of Via Padre Semeria is located on the southern side of Rome, along the Via Cristoforo Colombo (Catalano et al 2015) and close to the Aurelian walls. Land use was related to farming activities, as evidenced by the discovery of the ruins of a “*villa rustica*” and some hydraulic works (Ramieri, 1992), as well as analysis of skeletal stress markers suggesting that females were also involved in agricultural activity (Caldarini et al., 2015).

The baseline for the terrestrial protein component of the diet was set using 17 coeval faunal remains recovered from excavations at Rome (6 from Castel Malnome, 2 from Via Padre Semeria and 9 coming from Colosseum Area), to be used as ecological reference data, supplemented by previously published data for the same geographic and chronological frames. These published data were downloaded from IsoArcH database in several queries performed on or before October 30th, 2019 (Salesse et al., 2018; Prowse et al., 2001; O’Connel et al., 2019).

### ANALYTICAL METHODS

The extraction of collagen was individually performed following Longin’s protocol modified by Brown and colleagues (1988), which was also simultaneously applied to a modern bovine sample as a reference. In order to obtain a satisfactory yield of collagen, the extraction was performed on about 500 mg of bone powder collected by drilling the bones. The ultrafiltration step was also performed for all the samples in order to magnify the collagen concentration through >30 kDa Amicon® Ultra-4 Centrifugal Filter Units with Ultracel® membranes.

Each sample of collagen extract weighed 0.8-1.2 mg and was analyzed using an elemental analyzer isotope ratio mass spectrometer at the iCONa Laboratory of the University of Campania.

To test reliability and exclude contamination from exogenous carbon and nitrogen sources, the samples were compared against established criteria to ascertain the percentages of carbon and nitrogen, atomic C/N ratios, and collagen yields (Ambrose, 1990; DeNiro, 1985; van Klinken, 1999). Analytical precision was ± 0.3‰ for δ15N, reported with respect to AIR, and ± 0.1‰ for δ13C, reported with respect to the VPDB standard. Criteria for assessing the quality of preservation were carbon content, nitrogen content, and the C/N ratio (De Niro 1985; Ambrose and Norr,1993; van Klinken 1999).

Descriptive statistics and comparison tests were performed by R v.3.6.1 (R Core Team, 2017) The suggestions provided by Fraser and colleagues (2013) and recently further developed by Fontanals-Coll and colleagues (2016) were employed to detect the consumer’s role for humans compared to the available ecological resources. As described by the authors, this model uses the midpoint and the offsets between consecutive trophic levels to identify the effect of predators on their prey. Thus, the information based on faunal remains was organized according to typology (herbivores, omnivores, marine resources, freshwater organisms) and human data were plotted together in order to detect dietary preferences.

## RESULTS

The collagen extraction was performed for the whole sample but the preservation status of the extracted collagen led us to exclude some individual data: carbon content greater than or equal to 30%, nitrogen content greater than or equal to 10% (Ambrose 1990), and an atomic C/N ratio between 2.9 and 3.6 (DeNiro 1985) were the leading determinants for assessing suitable data. CM3 was depleted in elemental compositions but its C/N ratio and the associated δ^13^C and δ^15^ results are consistent with conspecific samples.

The extraction yield was not used as a criterion (Ambrose 1990) because the ultrafiltration technique was used. Only samples with yield of 0% were ruled out.

Faunal remains yielded enough collagen to be analyzed. Three bones of *Canis sp.* and two deer samples were recruited in Castel Malnome along with a cattle fragment and two herbivore fragments (sheep and cattle) from the Via Padre Semeria archaeological survey. The Colosseum Area domestics (one bird, one chicken, three pigs, two lambs, one hare and one cattle) return valid values too (Table 2).

**Table 2:** individual results for faunal remains.

| **Labcode** | **Species** | **%C** | **%N** | **C/N** | **δ<sup>13</sup>C‰** | **δ<sup>15</sup>N‰** |
| --- | --- | --- | --- | --- | --- | --- |
| **CM1** | *Dog* | 35.7 | 12.7 | 3.3 | -19.42 | 10.07 |
| **CM2** | *Dog* | 31.6 | 11.3 | 3.3 | -20.02 | 9.11 |
| **CM3** | *Dog* | 17.4 | 6.1 | 3.3 | -19.96 | 10.6 |
| **CM4** | *Deer* | 49.8 | 17.1 | 3.4 | -22.22 | 5.93 |
| **CM5** | *Deer* | 38.4 | 13.9 | 3.2 | -19.73 | 5.03 |
| **CM6** | *Cattle* | 39.8 | 13.6 | 3.4 | -21.52 | 5.28 |
| **PS1** | *Sheep* | 44 | 15.7 | 3.3 | -21.09 | 6.72 |
| **PS2** | *Cattle* | 45.5 | 16.3 | 3.3 | -20.07 | 6.73 |
| **COL1** | *Pig* | 45.5 | 16.6 | 3.2 | -20.44 | 6.76 |
| **COL2** | *Goat* | 84.6 | 31.3 | 3.2 | -20.61 | 4.24 |
| **COL3** | *Pig* | 41.6 | 15.3 | 3.2 | -20.24 | 4.58 |
| **COL4** | *Chicken* | 50 | 18.5 | 3.2 | -20.81 | 5.46 |
| **COL5** | *Bird* | 31.4 | 11.2 | 3.3 | -19.93 | 8.14 |
| **COL6** | *Sheep* | 42.4 | 15 | 3.3 | -21.82 | 5.42 |
| **COL7** | *Cattle* | 48 | 17.3 | 3.2 | -20.81 | 5.48 |
| **COL8** | *Hare* | 31.1 | 11.4 | 3.2 | -21.93 | 3.82 |
| **COL9** | *Pig* | 33.2 | 11.8 | 3.3 | -20.03 | 6.33 |

The obtained faunal δ^13^C values are consistent with a C_3_ European ecosystem (Schwarcz and Schoeninger 1991) and the δ^15^N signature suggests the proper trophic level for the identified species.

Out of 214 human samples, only 199 fit the quality criteria. Considering all 199 human individuals, δ^13^C ranges from −19,95‰ to −14,78‰, whereas δ^15^N values are between 7,17‰ and 10,00‰ (Table 3).

**Table 3:** Individual results for humans. F: female; M: male; Ind: gender not available.

| Site | Sample | Sex | Age | %C | %N | C/N | $\delta^{13}\text{C}\text{‰}$ | $\delta^{15}\text{N}\text{‰}$ |
| --- | --- | --- | --- | --- | --- | --- | --- | --- |
| CM | CM1 | M | 50-x | 39.9 | 13.9 | 3.3 | -18.71 | 11.18 |
| CM | CM2 | M | 40-46 | 42 | 15.1 | 3.2 | -18.72 | 11.38 |
| CM | CM3 | M | 40-49 | 41.6 | 14.8 | 3.3 | -19.1 | 11.37 |
| CM | CM4 | M | 30-39 | 40.8 | 14.6 | 3.3 | -18.74 | 12.24 |
| CM | CM5 | M | 20-29 | 39.6 | 15.1 | 3.1 | -19 | 9.8 |
| CM | CM6 | F | 20-29 | 43.7 | 15.2 | 3.4 | -19.31 | 10.62 |
| CM | CM7 | F | 20-29 | 40.2 | 15.2 | 3.1 | -19.31 | 9.34 |
| CM | CM8 | M | 40-49 | 40.1 | 15.3 | 3.1 | -19.47 | 10.18 |
| CM | CM9 | M | 40-49 | 40.1 | 15.2 | 3.1 | -19.76 | 9.19 |
| CM | CM10 | M | 20-29 | 38.9 | 15.1 | 3 | -19.4 | 9.68 |
| CM | CM11 | M | 40-49 | 38.1 | 12.3 | 3.6 | -19.12 | 11.21 |
| CM | CM13 | M | 50-x | 40.1 | 14.8 | 3.2 | -19.09 | 10.98 |
| CM | CM14 | M | 40-49 | 40.2 | 13.7 | 3.4 | -19.01 | 10.54 |
| CM | CM15 | M | 40-49 | 36.1 | 12.4 | 3.4 | -19.43 | 11.76 |
| CM | CM16 | M | 20-29 | 32.2 | 11 | 3.4 | -20.38 | 7.17 |
| CM | CM17 | M | 20-29 | 34.4 | 13.9 | 2.9 | -18.88 | 11.3 |
| CM | CM18 | M | 40-49 | 39.5 | 13.7 | 3.4 | -19.59 | 11.1 |
| CM | CM19 | M | 30-39 | 40.7 | 14.2 | 3.3 | -19.03 | 11.49 |
| CM | CM20 | Ind | 13-19 | 42.1 | 13.6 | 3.6 | -20.57 | 8.47 |
| CM | CM21 | F | 13-19 | 41.4 | 13.3 | 3.6 | -20.81 | 7.65 |
| CM | CM22 | M | 40-49 | 41.3 | 14.1 | 3.4 | -18.79 | 11.05 |
| CM | CM23 | F | 30-39 | 41.2 | 15.1 | 3.2 | -20.42 | 8.00 |
| CM | CM24 | M | 40-49 | 45.3 | 15.6 | 3.4 | -18.8 | 12.31 |
| CM | CM25 | M | 30-39 | 44.5 | 15 | 3.5 | -19.00 | 9.70 |
| CM | CM26 | M | 40-49 | 44.6 | 15.6 | 3.3 | -19.22 | 11.17 |
| CM | CM27 | M | 20-29 | 44.6 | 15.5 | 3.4 | -18.7 | 8.98 |
| CM | CM28 | M | 40-49 | 44 | 15.3 | 3.4 | -19.13 | 12.48 |
| CM | CM29 | M | 20-29 | 45.4 | 15.4 | 3.4 | -18.89 | 12.61 |
| CM | CM30 | M | 40-49 | 45 | 15.7 | 3.3 | -19.45 | 10.34 |
| CM | CM31 | F | 40-49 | 43.4 | 15.1 | 3.4 | -18.96 | 11.47 |
| CM | CM32 | M | 50-x | 41.9 | 14.7 | 3.3 | -18.74 | 11.91 |
| CM | CM33 | M | 40-49 | 40.1 | 14.8 | 3.2 | -20.39 | 8.54 |
| CM | CM34 | M | 20-29 | 40.3 | 14.6 | 3.2 | -14.78 | 11.04 |
| CM | CM35 | Ind | 13-19 | 40.2 | 14.7 | 3.2 | -19.36 | 11.23 |
| CM | CM36 | Ind | 13-19 | 41.2 | 14.3 | 3.4 | -19.07 | 11.25 |
| CM | CM37 | F | 30-39 | 42.1 | 13.9 | 3.5 | -19.24 | 11.68 |
| CM | CM39 | F | 20-29 | 42.3 | 14.8 | 3.3 | -19.24 | 12.03 |
| CM | CM40 | M | 40-49 | 44.7 | 15.7 | 3.3 | -19.57 | 11.03 |
| CM | CM41 | M | 40-49 | 42.3 | 15.1 | 3.3 | -20.38 | 9.09 |
| CM | CM42 | F | 20-29 | 40.6 | 15.9 | 3 | -19.09 | 10.54 |
| CM | CM43 | M | 30-39 | 42.3 | 15.8 | 3.1 | -19.16 | 10.59 |
| CM | CM47 | M | 20-29 | 43.2 | 14.2 | 3.5 | -19.01 | 12.92 |
| CM | CM48 | F | 40-49 | 41.1 | 14.9 | 3.2 | -19.15 | 9.72 |
| CM | CM49 | M | 30-39 | 41.2 | 14.8 | 3.2 | -18.93 | 10.5 |
| CM | CM50 | M | 40-49 | 41.5 | 14.3 | 3.4 | -19.06 | 10.97 |
| CM | CM51 | Ind | 13-19 | 41.6 | 13.9 | 3.5 | -19.55 | 11.51 |
| CM | CM52 | M | 50-x | 43.9 | 15.5 | 3.3 | -17.02 | 12.44 |
| CM | CM53 | M | 30-39 | 42.4 | 14.2 | 3.5 | -19.77 | 9.19 |
| CM | CM54 | F | 30-39 | 40.8 | 14.8 | 3.2 | -19.14 | 11.74 |
| CM | CM55 | M | 30-39 | 41.9 | 15.1 | 3.2 | -18.21 | 12.57 |
| CM | CM56 | Ind. | >21 | 42.1 | 14.1 | 3.5 | -19.15 | 10.29 |
| CM | CM57 | M | 20-29 | 42.9 | 14.2 | 3.5 | -19.11 | 9.68 |
| CM | CM58 | F | 20-29 | 42.7 | 13.9 | 3.6 | -19.27 | 11.72 |
| CM | CM60 | F | 50-x | 38.8 | 13.8 | 3.3 | -19.05 | 10.93 |
| CM | CM61 | M | 30-39 | 39.8 | 14.2 | 3.3 | -19.54 | 11.29 |
| CM | CM62 | M | 30-39 | 39.9 | 14.4 | 3.2 | -19.32 | 11.98 |
| CM | CM63 | F | 30-39 | 41.1 | 14.6 | 3.3 | -19.34 | 11.34 |
| CM | CM64 | F | 40-49 | 42.4 | 14.8 | 3.3 | -18.91 | 11.01 |
| CM | CM65 | M | 40-49 | 40.8 | 15.1 | 3.2 | -19.78 | 10.61 |
| CM | CM66 | M | 30-39 | 41.8 | 15.3 | 3.2 | -20.64 | 9.31 |
| CM | CM67 | M | 40-49 | 42.2 | 15 | 3.3 | -19.36 | 11.6 |
| CM | CM68 | F | 40-49 | 39.8 | 15 | 3.1 | -19.12 | 11.97 |
| CM | CM69 | M | 40-49 | 40 | 15.3 | 3.1 | -20.09 | 12.16 |
| CM | CM70 | Ind | 13-19 | 40.8 | 15.1 | 3.2 | -19.49 | 11.67 |
| CM | CM71 | M | 30-39 | 40.6 | 15 | 3.2 | -18.84 | 9.76 |
| CM | CM73 | F | 20-29 | 41 | 14.3 | 3.3 | -18.82 | 11.44 |
| CM | CM74 | F | 20-29 | 43.8 | 14.8 | 3.5 | -19.33 | 11.43 |
| CM | CM75 | M | 40-49 | 43.5 | 14.4 | 3.5 | -19.76 | 9.26 |
| CM | CM76 | M | 30-39 | 41 | 14.6 | 3.3 | -18.73 | 10.9 |
| CM | CM77 | M | 30-39 | 41.6 | 14.1 | 3.4 | -19.89 | 10.31 |
| CM | CM78 | F | 30-39 | 44.6 | 15.7 | 3.3 | -19.21 | 11.67 |
| CM | CM79 | F | 30-39 | 43.9 | 14.9 | 3.4 | -19.29 | 11.8 |
| PS | PS1 | M | 20-29 | 43.7 | 15.5 | 3.3 | -18.86 | 11.86 |
| PS | PS4 | M | 13-19 | 42.2 | 14.5 | 3.4 | -19.68 | 10.27 |
| PS | PS5 | M | 30-39 | 31.6 | 10.8 | 3.4 | -18.94 | 10.35 |
| PS | PS6 | F | 20-29 | 39.4 | 13.9 | 3.3 | -19.41 | 10.65 |
| PS | PS7 | F | 40-49 | 43.7 | 15.2 | 3.4 | -18.26 | 12.43 |
| PS | PS8 | F | 13-19 | 40.9 | 14.3 | 3.3 | -19.00 | 10.47 |
| PS | PS9 | M | 30-39 | 40.1 | 15.1 | 3.1 | -19.31 | 11.43 |
| PS | PS10 | M | 20-29 | 44.7 | 16.2 | 3.2 | -18.9 | 11.99 |
| PS | PS11 | M | 30-39 | 44.4 | 15.3 | 3.4 | -19.32 | 12.1 |
| PS | PS12 | F | 40-49 | 38.6 | 13.9 | 3.2 | -19.5 | 10.67 |
| PS | PS13 | F | 20-29 | 44.3 | 14.7 | 3.5 | -19.43 | 10.86 |
| PS | PS14 | F | 20-29 | 39.7 | 14 | 3.3 | -18.99 | 11.34 |
| PS | PS15 | F | 30-39 | 43.1 | 15.5 | 3.2 | -19.05 | 10.00 |
| PS | PS16 | F | 20-29 | 33.8 | 11.3 | 3.5 | -19.95 | 11.3 |
| PS | PS17 | M | 40-49 | 40.9 | 14.2 | 3.4 | -19.24 | 11.33 |
| PS | PS18 | Ind | 07-12 | 37.3 | 13 | 3.3 | -19.34 | 11.77 |
| PS | PS19 | F | 20-29 | 40.9 | 14.3 | 3.3 | -19.35 | 11.72 |
| PS | PS20 | F | 20-29 | 26.5 | 8.7 | 3.6 | -19.76 | 10.88 |
| PS | PS21 | F | 20-29 | 23.8 | 7.7 | 3.6 | -19.57 | 11.41 |
| PS | PS22 | M | 40-49 | 24.9 | 8.3 | 3.5 | -19.51 | 10.01 |
| PS | PS23 | M | 20-29 | 39.5 | 14.2 | 3.2 | -18.61 | 11.43 |
| PS | PS24 | M | 40-49 | 41.8 | 15 | 3.3 | -18.58 | 12.37 |
| PS | PS25 | F | 13-19 | 25.5 | 8.5 | 3.5 | -18.58 | 13.23 |
| PS | PS26 | M | 20-29 | 41.6 | 14.6 | 3.3 | -19.25 | 10.45 |
| PS | PS27 | F | 20-29 | 40.8 | 14.2 | 3.4 | -19.00 | 11.79 |
| PS | PS28 | F | 20-29 | 36 | 12.3 | 3.4 | -19.11 | 12.06 |
| PS | PS30 | M | 20-29 | 49.6 | 17.7 | 3.3 | -18.05 | 13.05 |
| QCP | QCP1 | F | 30-39 | 44.4 | 15 | 3.5 | -19.52 | 9.93 |
| QCP | QCP2 | M | 40-49 | 44.1 | 14.8 | 3.5 | -18.58 | 11.53 |
| QCP | QCP3 | Ind | 0-6 | 50.8 | 18.5 | 3.2 | -19.12 | 8.78 |
| QCP | QCP5 | Ind | 07-12 | 43.3 | 15.1 | 3.3 | -19.38 | 8.48 |
| QCP | QCP6 | M | 40-49 | 43.7 | 15.9 | 3.2 | -18.70 | 12.31 |
| QCP | QCP7 | Ind | 0-6 | 36.7 | 12.9 | 3.3 | -19.70 | 9.13 |
| QCP | QCP8 | M | 30-39 | 41.7 | 15.2 | 3.2 | -19.32 | 9.74 |
| QCP | QCP9 | M | 30-39 | 43.2 | 15.5 | 3.3 | -18.83 | 10.23 |
| QCP | QCP10 | Ind | 0-6 | 42.5 | 15.5 | 3.2 | -19.2 | 9.00 |
| QCP | QCP11 | F | 20-29 | 37 | 13.2 | 3.3 | -19.51 | 8.74 |
| QCP | QCP12 | F | 20-29 | 38.8 | 15.2 | 3 | -18.76 | 9.58 |
| QCP | QCP13 | Ind | 0-6 | 39.7 | 14.1 | 3.3 | -19.42 | 8.84 |
| QCP | QCP14 | F | 30-39 | 42.2 | 15.2 | 3.2 | -19.05 | 9.04 |
| QCP | QCP15 | Ind | 0-6 | 46.5 | 16.9 | 3.2 | -19.24 | 9.35 |
| QCP | QCP16 | Ind | 0-6 | 44.5 | 16.1 | 3.2 | -18.50 | 10.85 |
| QCP | QCP17 | M | 20-29 | 43.1 | 14.8 | 3.4 | -19.83 | 8.08 |
| QCP | QCP18 | Ind | 0-6 | 40.8 | 14.6 | 3.3 | -18.94 | 10.12 |
| QCP | QCP19 | Ind | 0-6 | 43.1 | 15 | 3.4 | -19.47 | 11.76 |
| QCP | QCP20 | Ind | 0-6 | 40.1 | 15.1 | 3.1 | -19.37 | 12.14 |
| QCP | QCP21 | F | 30-39 | 44.4 | 16.2 | 3.2 | -20.53 | 8.39 |
| QCP | QCP22 | M | 30-39 | 42.2 | 14.6 | 3.4 | -19.42 | 8.75 |
| QCP | QCP23 | Ind | 0-6 | 31 | 10.9 | 3.3 | -19.72 | 8.33 |
| QCP | QCP24 | Ind | 0-6 | 35.6 | 12.5 | 3.3 | -19.13 | 13.52 |
| QCP | QCP25 | Ind | 0-6 | 36.8 | 13 | 3.3 | -18.35 | 14.30 |
| QCP | QCP27 | Ind | 0-6 | 40.3 | 13.9 | 3.4 | -20.07 | 9.66 |
| QCP | QCP28 | M | 20-29 | 40.7 | 14.3 | 3.3 | -19.58 | 7.72 |
| QCP | QCP29 | F | 40-49 | 39.2 | 13.8 | 3.3 | -19.81 | 7.69 |
| QCP | QCP30 | Ind | 0-6 | 43.2 | 15.7 | 3.2 | -18.71 | 11.79 |
| QCP | QCP31 | Ind | 13-19 | 39.2 | 14.1 | 3.2 | -19.46 | 9.70 |
| QCP | QCP32 | M | 40-49 | 44 | 14.9 | 3.4 | -20.01 | 9.11 |
| QCP | QCP33 | F | 30-39 | 37 | 13.5 | 3.2 | -19.08 | 10.31 |
| QCP | QCP34 | Ind | 0-6 | 53.3 | 19.2 | 3.2 | -18.96 | 11.37 |
| QCP | QCP35 | M | 40-49 | 34.7 | 11.9 | 3.4 | -19.28 | 10.16 |
| QCP | QCP36 | Ind | 0-6 | 38.8 | 14.9 | 3 | -19.90 | 11.72 |
| QCP | QCP37 | Ind | 07-12 | 41.5 | 15.1 | 3.2 | -18.98 | 10.82 |
| QCP | QCP39 | M | 13-19 | 42.2 | 15.5 | 3.2 | -18.17 | 8.84 |
| QCP | QCP40 | M | 30-39 | 37.7 | 13.7 | 3.2 | -18.87 | 11.39 |
| CBN | CBN1 | M | 30-39 | 36.6 | 12.8 | 3.3 | -16.49 | 12.1 |
| **CBN** | CBN2 | M | 50-x | 32.7 | 11.5 | 3.3 | -18.20 | 11.1 |
| **CBN** | CBN3 | M | 20-29 | 39.9 | 13.4 | 3.5 | -20.01 | 9.29 |
| **CBN** | CBN4 | F | 20-29 | 38.2 | 13.4 | 3.3 | -19.71 | 8.59 |
| **CBN** | CBN5 | M | 40-49 | 25.4 | 8.6 | 3.4 | -18.66 | 11.3 |
| **CBN** | CBN6 | F | 30-39 | 41.4 | 13.9 | 3.5 | -18.90 | 11.94 |
| **CBN** | CBN7 | M | 20-29 | 30.1 | 10.2 | 3.4 | -20.37 | 11.81 |
| **CBN** | CBN8 | M | 13-19 | 38.7 | 13.5 | 3.3 | -19.19 | 11.48 |
| **CBN** | CBN9 | Ind | 13-19 | 43.9 | 15.2 | 3.4 | -18.49 | 11.25 |
| **CBN** | CBN10 | M | 30-39 | 30 | 9.8 | 3.6 | -19.00 | 11.57 |
| **CBN** | CBN11 | Ind | 13-19 | 27.1 | 8.7 | 3.6 | -18.6 | 11.28 |
| **CBN** | CBN12 | Ind | 0-6 | 32.8 | 11.2 | 3.4 | -19.2 | 11.41 |
| **CBN** | CBN13 | Ind | 13-19 | 37.7 | 13.1 | 3.4 | -19.14 | 10.57 |
| **CBN** | CBN14 | M | 40-49 | 41.7 | 14.6 | 3.3 | -18.55 | 12.04 |
| **CBN** | CBN15 | M | 20-29 | 26.2 | 8.9 | 3.4 | -18.96 | 12.37 |
| **CBN** | CBN16 | M | 30-39 | 44 | 15.2 | 3.4 | -19.04 | 11.89 |
| **CBN** | CBN18 | F | 30-39 | 45 | 15.8 | 3.3 | -19.64 | 8.36 |
| **CBN** | CBN19 | Ind | 13-19 | 35.9 | 12.1 | 3.5 | -18.70 | 11.39 |
| **CBM** | CBM1 | M | 30-39 | 41.9 | 15.2 | 3.2 | -18.34 | 12.22 |
| **CBM** | CBM2 | F | 20-29 | 46.2 | 16.5 | 3.3 | -18.58 | 11.91 |
| **CBM** | CBM3 | Ind | 13-19 | 41 | 14.4 | 3.3 | -18.78 | 11.59 |
| **CBM** | CBM4 | M | 30-39 | 42.1 | 14.7 | 3.3 | -18.22 | 11.84 |
| **CBM** | CBM5 | M | 40-49 | 48.7 | 15.6 | 3.6 | -18.71 | 11.37 |
| **CBM** | CBM6 | F | 40-49 | 43.2 | 15.3 | 3.3 | -18.92 | 11.3 |
| **CBM** | CBM7 | F | >20 | 40.5 | 14.2 | 3.3 | -18.94 | 11.61 |
| **CBM** | CBM8 | Ind | 07-12 | 41.3 | 14.4 | 3.3 | -18.61 | 11.08 |
| **CBM** | CBM9 | F | 30-39 | 39.3 | 13.7 | 3.3 | -18.63 | 11.26 |
| **CBM** | CBM10 | Ind | 13-19 | 43.7 | 15.2 | 3.4 | -19.13 | 10.81 |
| **CBM** | CBM11 | Ind | 07-12 | 45.6 | 15.8 | 3.4 | -18.90 | 9.72 |
| **CBM** | CBM12 | Ind | 13-19 | 43.5 | 15.3 | 3.3 | -18.96 | 10.66 |
| **CBM** | CBM13 | Ind | 07-12 | 44.2 | 15.6 | 3.3 | -19.12 | 10.62 |
| **CBM** | CBM14 | M | 20-29 | 41.9 | 14.7 | 3.3 | -18.74 | 11.02 |
| **CBM** | CBM15 | F | 20-29 | 42.3 | 14.9 | 3.3 | -19.26 | 9.62 |
| **CBM** | CBM16 | M | 50-x | 46.9 | 16.6 | 3.3 | -18.89 | 11.49 |
| **CBM** | CBM17 | F | 30-39 | 46.3 | 16.2 | 3.3 | -19.27 | 10.53 |
| **CBM** | CBM18 | Ind | 07-12 | 43.6 | 15.6 | 3.3 | -18.83 | 11.6 |
| **CBM** | CBM19 | Ind | 07-12 | 41.3 | 14.5 | 3.3 | -19.30 | 11.78 |
| **CBM** | CBM20 | Ind | 07-12 | 40.3 | 14.2 | 3.3 | -19.12 | 8.58 |
| **CBM** | CBM21 | M | 13-19 | 41.6 | 14.6 | 3.3 | -19.09 | 11.03 |
| **CBM** | CBM22 | Ind | 07-12 | 46.1 | 16.2 | 3.3 | -19.20 | 9.77 |
| **CBM** | CBM23 | Ind | 07-12 | 41.9 | 14.7 | 3.3 | -20.38 | 8.13 |
| **CBM** | CBM24 | M | 40-49 | 47.3 | 17.2 | 3.2 | -18.41 | 12.2 |
| **CBM** | CBM25 | Ind | 13-19 | 40.6 | 14.3 | 3.3 | -19.70 | 8.59 |
| **CBM** | CBM26 | Ind | 07-12 | 44.4 | 15.7 | 3.3 | -19.19 | 10.06 |
| **CBQ** | CBQ1 | M | 50-X | 44.7 | 14.9 | 3.5 | -18.96 | 12.57 |
| **CBQ** | CBQ2 | M | 13-19 | 44.7 | 16.2 | 3.2 | -18.93 | 10.65 |
| **CBQ** | CBQ3 | F | 13-19 | 33.2 | 11.4 | 3.4 | -20.24 | 8.33 |
| **CBQ** | CBQ4 | M | >20 | 39.8 | 13.8 | 3.4 | -19.25 | 10.63 |
| **CBQ** | CBQ5 | M | 50-X | 14.6 | 4.9 | 3.5 | -19.27 | 10.25 |
| **CBQ** | CBQ6 | Ind | 07-12 | 36.4 | 13.3 | 3.2 | -18.90 | 11.28 |
| **CBQ** | CBQ7 | Ind | 0-6 | 47.1 | 16.8 | 3.3 | -18.83 | 10.74 |
| **CBQ** | CBQ8 | Ind | 0-6 | 43 | 15.3 | 3.3 | -19.72 | 10.74 |
| **CBQ** | CBQ9 | F | 20-29 | 34.4 | 11 | 3.6 | -19.45 | 11.44 |
| **CBQ** | CBQ10 | Ind | 0-6 | 42.6 | 15.1 | 3.3 | -18.95 | 11.92 |
| **CBQ** | CBQ11 | M | 40-49 | 42.2 | 15.1 | 3.3 | -18.78 | 11.97 |
| **CBQ** | CBQ12 | M | 50-X | 42 | 14.7 | 3.3 | -19.22 | 9.43 |
| **CBQ** | CBQ13 | F | 40-49 | 39.6 | 13.8 | 3.3 | -17.62 | 12.63 |
| **CBQ** | CBQ14 | Ind | 0-6 | 39.9 | 14.8 | 3.1 | -19.21 | 11.7 |
| **CBQ** | CBQ15 | F | 50-X | 40.2 | 14.4 | 3.3 | -18.59 | 11.14 |
| **CBQ** | CBQ16 | M | 40-49 | 42.8 | 15.1 | 3.3 | -18.66 | 9.60 |
| **CBQ** | CBQ17 | F | 30-39 | 42.9 | 15 | 3.3 | -19.17 | 10.77 |
| **CBQ** | CBQ18 | Ind | 0-6 | 33.5 | 11.7 | 3.3 | -19.75 | 11.15 |
| **CBQ** | CBQ19 | F | >20 | 41.5 | 14.7 | 3.3 | -19.13 | 12.56 |

The overall data distribution and the density plots indicate a certain heterogeneity: δ^13^C and δ^15^N values range between 1.90‰ and 6.03‰ and between 3.23‰ and 6.61‰ respectively (Figure 2).

**Figure 2:**
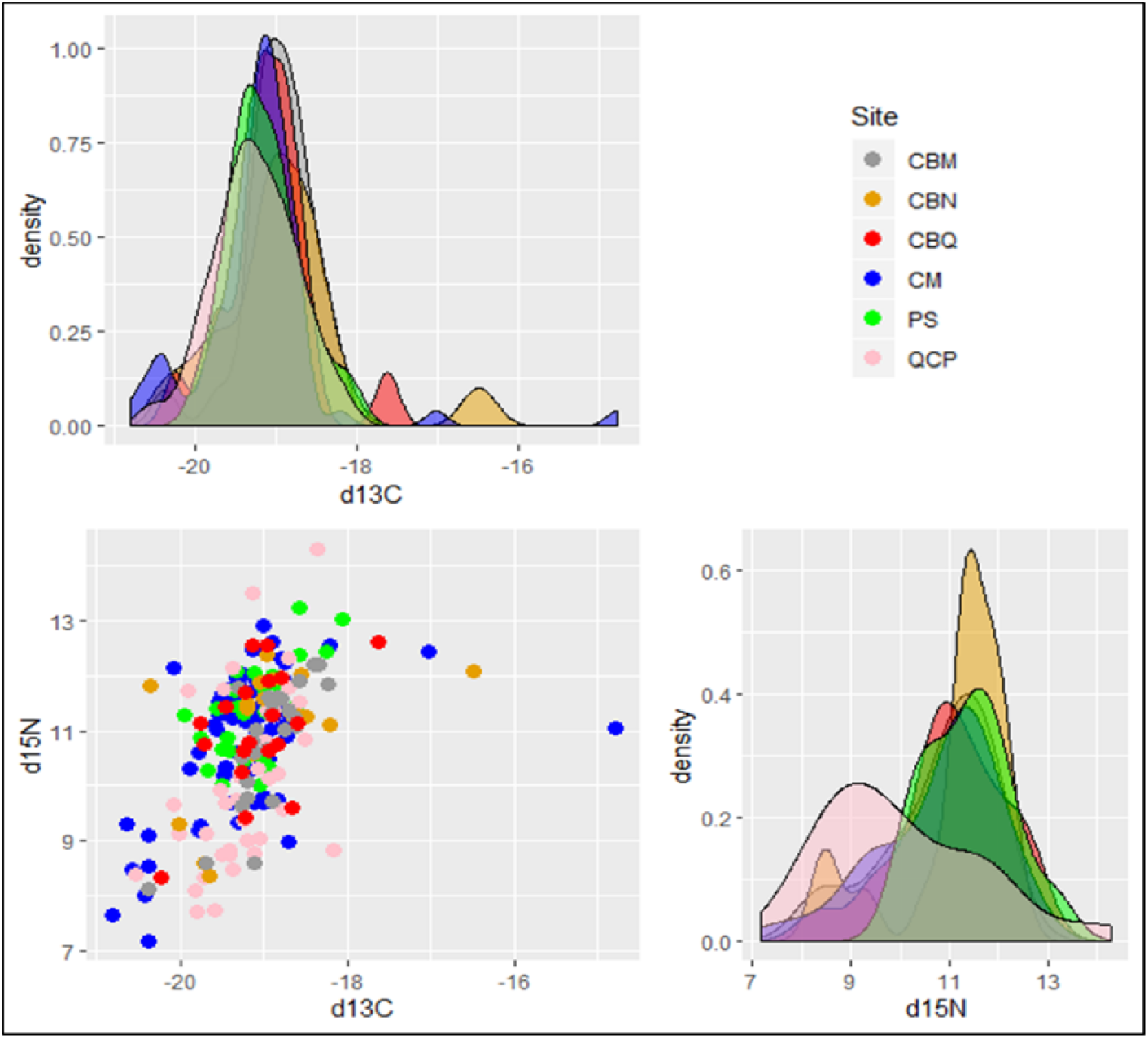
Plot for δ13C than δ15N values. Density plots refer to each axis.

Indeed, the wider range for δ^13^C than δ^15^N could be due to the presence of a few “positive” outliers such as CM34 and CM52 (−14.80‰ and −17.00‰ for δ^13^C) as well as CBN1(−16.5‰ for δ^13^C) and CBQ13 (−17.60‰ for δ^13^C). Likewise, some lower-δ^15^N outliers in CBN (CBN3, CBN4, and CBN18 with 9.29‰, 8.59‰, and 8.36‰ respectively) and the spanned values for QCP samples account for the wide range detected for δ^15^N.

The values are consistent with an overall diet mainly based on terrestrial resources. All the human samples rely on a higher trophic level than the primary consumer faunal samples, with no clear indication of exclusive marine food source consumption, although appreciable consumption of these cannot be ruled out, especially for some people at Castel Malnome and Casal Bertone, both at the necropolis and the mausoleum, due to the less negative δ^13^C data.

The sample stratification according to the necropolis could allow us to evaluate putative differences in food source exploitation. The descriptive statistics for δ^13^C and δ^15^N for the six funerary areas were calculated (Table 4).

**Table 4:** Descriptive statistics for the 6 necropolises. These statistics are the minimal value (min), the maximal value (max), the range (range, that is, max-min), the median (median), the mean (mean), the standard error on the mean (SE.mean), the confidence interval of the mean (CI.mean) at the p=0.95 level, the variance (var), the standard deviation (std.dev) and the variation coefficient (coef.var) defined as the standard deviation divided by the mean.

|  | **CBM** |  | **CBN** |  | **CBQ** |  | **CM** |  | **PS** |  | **QCP** |  |
| --- | --- | --- | --- | --- | --- | --- | --- | --- | --- | --- | --- | --- |
| **Sample size** | 26 |  | 18 |  | 19 |  | 72 |  | 27 |  | 37 |  |
| **Average C:N** | 3.3 |  | 3.4 |  | 3.3 |  | 3.3 |  | 3.4 |  | 3.3 |  |
| | $\delta^{13}\text{C}$ | $\delta^{15}\text{N}$ | $\delta^{13}\text{C}$ | $\delta^{15}\text{N}$ | $\delta^{13}\text{C}$ | $\delta^{15}\text{N}$ | $\delta^{13}\text{C}$ | $\delta^{15}\text{N}$ | $\delta^{13}\text{C}$ | $\delta^{15}\text{N}$ | $\delta^{13}\text{C}$ | $\delta^{15}\text{N}$ |
| **Min** | -20.38 | 8.13 | -20.37 | 8.36 | -20.24 | 8.33 | -20.81 | 7.17 | -19.95 | 10.00 | -20.53 | 7.69 |
| **Max** | -18.22 | 12.22 | -16.49 | 12.37 | -17.62 | 12.63 | -14.78 | 12.92 | -18.05 | 13.23 | -18.17 | 14.30 |
| **Range** | 2.16 | 4.09 | 3.88 | 4.01 | 2.62 | 4.30 | 6.03 | 5.75 | 1.90 | 3.23 | 2.36 | 6.61 |
| **median** | -18.93 | 11.06 | -18.98 | 11.40 | -19.13 | 11.14 | -19.16 | 11.08 | -19.24 | 11.41 | -19.28 | 9.70 |
| **mean** | -18.97 | 10.78 | -18.94 | 11.10 | -19.09 | 11.03 | -19.22 | 10.80 | -19.13 | 11.38 | -19.26 | 10.03 |
| **SE.mean** | 0.09 | 0.22 | 0.19 | 0.28 | 0.12 | 0.26 | 0.09 | 0.15 | 0.09 | 0.17 | 0.08 | 0.26 |
| **CI.mean** | 0.18 | 0.46 | 0.41 | 0.58 | 0.26 | 0.54 | 0.18 | 0.29 | 0.18 | 0.34 | 0.17 | 0.53 |
| **var** | 0.19 | 1.27 | 0.68 | 1.37 | 0.29 | 1.26 | 0.62 | 1.52 | 0.20 | 0.76 | 0.26 | 2.54 |
| **std.dev** | 0.44 | 1.13 | 0.83 | 1.17 | 0.54 | 1.12 | 0.78 | 1.23 | 0.45 | 0.87 | 0.51 | 1.59 |
| **coef.var** | -0.02 | 0.10 | -0.04 | 0.11 | -0.03 | 0.10 | -0.04 | 0.11 | -0.02 | 0.08 | -0.03 | 0.16 |

The osteological evaluation of the human remains allowed us to determine the gender of all individuals, which led us to dissect the variability in food consumption between males and females as summarized in Table 5.

**Table 5:** Basic descriptive statistics for the 6 necropolises stratified according to genders.

|  | CBM |  | CBN |  | CBQ |  | CM |  | PS |  | QCP |  |
| --- | --- | --- | --- | --- | --- | --- | --- | --- | --- | --- | --- | --- |
|  | Males (n=7) |  | Males (n=10) |  | Males (n=7) |  | Males (n=47) |  | Males (n=12) |  | Males (n=11) |  |
| | $\delta^{13}\text{C}$ | $\delta^{15}\text{N}$ | $\delta^{13}\text{C}$ | $\delta^{15}\text{N}$ | $\delta^{13}\text{C}$ | $\delta^{15}\text{N}$ | $\delta^{13}\text{C}$ | $\delta^{15}\text{N}$ | $\delta^{13}\text{C}$ | $\delta^{15}\text{N}$ | $\delta^{13}\text{C}$ | $\delta^{15}\text{N}$ |
| **Mean** | -18.63 | 11.6 | -18.85 | 11.49 | -19.01 | 10.73 | -19.14 | 10.78 | -19.02 | 11.39 | -19.14 | 9.8 |
| **Median** | -18.71 | 11.49 | -18.98 | 11.69 | -18.96 | 10.63 | -19.11 | 11.03 | -19.09 | 11.43 | -19.28 | 9.74 |
| **Variance** | 0.1 | 0.25 | 1.11 | 0.75 | 0.06 | 1.36 | 0.8 | 1.52 | 0.21 | 0.9 | 0.32 | 2.19 |
|  | Females (n=6) |  | Females (n=3) |  | Females (n=6) |  | Females (n=19) |  | Females (n=14) |  | Females (n=7) |  |
| | $\delta^{13}\text{C}$ | $\delta^{15}\text{N}$ | $\delta^{13}\text{C}$ | $\delta^{15}\text{N}$ | $\delta^{13}\text{C}$ | $\delta^{15}\text{N}$ | $\delta^{13}\text{C}$ | $\delta^{15}\text{N}$ | $\delta^{13}\text{C}$ | $\delta^{15}\text{N}$ | $\delta^{13}\text{C}$ | $\delta^{15}\text{N}$ |
| **Mean** | -18.93 | 11.04 | -19.42 | 9.63 | -19.03 | 11.14 | -19.31 | 10.85 | -19.21 | 11.34 | -19.47 | 9.1 |
| **Median** | -18.93 | 11.28 | -19.64 | 8.59 | -19.15 | 11.29 | -19.24 | 11.43 | -19.23 | 11.32 | -19.51 | 9.04 |
| **Variance** | 0.09 | 0.69 | 0.2 | 4.01 | 0.77 | 2.47 | 0.23 | 1.66 | 0.2 | 0.73 | 0.36 | 0.83 |

## DISCUSSION

### DATA INTEGRATION

Two previously analyzed samples from Casal Bertone necropolis and Casal Bertone mausoleum (Killgrove and Tykot, 2013, Table A.1) were appended to the presented data sets in order to create larger samples pertaining to 231 individuals, whose basic descriptive statistics are listed in Table 6.

**Table 6:** Basic descriptive statistics for the whole sample.

|  | CBM |  | CBN |  | CBQ |  | CM |  | PS |  | QCP |  |
| --- | --- | --- | --- | --- | --- | --- | --- | --- | --- | --- | --- | --- |
| Sample size | 38 |  | 38 |  | 19 |  | 72 |  | 27 |  | 37 |  |
| | $\delta^{13}\text{C}$ | $\delta^{15}\text{N}$ | $\delta^{13}\text{C}$ | $\delta^{15}\text{N}$ | $\delta^{13}\text{C}$ | $\delta^{15}\text{N}$ | $\delta^{13}\text{C}$ | $\delta^{15}\text{N}$ | $\delta^{13}\text{C}$ | $\delta^{15}\text{N}$ | $\delta^{13}\text{C}$ | $\delta^{15}\text{N}$ |
| min | -20.38 | 7.00 | -20.37 | 7.20 | -20.24 | 8.33 | -20.81 | 7.17 | -19.95 | 10.00 | -20.53 | 7.69 |
| max | -17.50 | 12.22 | -16.49 | 13.20 | -17.62 | 12.63 | -14.78 | 12.92 | -18.05 | 13.23 | -18.17 | 14.30 |
| range | 2.88 | 5.22 | 3.88 | 6.00 | 2.62 | 4.30 | 6.03 | 5.75 | 1.90 | 3.23 | 2.36 | 6.61 |
| median | -18.76 | 10.91 | -18.60 | 11.10 | -19.13 | 11.14 | -19.16 | 11.08 | -19.24 | 11.41 | -19.28 | 9.70 |
| mean | -18.73 | 10.49 | -18.60 | 10.64 | -19.09 | 11.03 | -19.22 | 10.80 | -19.13 | 11.38 | -19.26 | 10.03 |
| SE.mean | 0.09 | 0.21 | 0.13 | 0.23 | 0.12 | 0.26 | 0.09 | 0.15 | 0.09 | 0.17 | 0.08 | 0.26 |
| CI.mean | 0.19 | 0.43 | 0.27 | 0.47 | 0.26 | 0.54 | 0.18 | 0.29 | 0.18 | 0.34 | 0.17 | 0.53 |
| var | 0.34 | 1.75 | 0.67 | 2.03 | 0.29 | 1.26 | 0.62 | 1.52 | 0.20 | 0.76 | 0.26 | 2.54 |
| std.dev | 0.58 | 1.32 | 0.82 | 1.43 | 0.54 | 1.12 | 0.78 | 1.23 | 0.45 | 0.87 | 0.51 | 1.59 |
| coef.var | -0.03 | 0.13 | -0.04 | 0.13 | -0.03 | 0.10 | -0.04 | 0.11 | -0.02 | 0.08 | -0.03 | 0.16 |

Furthermore, we are aware that the very restricted sample size for the faunal remains (17 animals) might be only minimally useful for representing the animal baseline, resulting in a bias for the dietary reconstruction of this large urban area. Thus, coeval data was collected by IsoArch Database and from the literature (O’Connel et al., 2019) in order to solve this issue. The faunal remains of 48 animals from several species (Table A.2) made up the whole sample used to support this data set as local ecological reference data for Imperial Rome.

A few diachronic samples (mid-5^th^–early-6^th^ centuries CE) were included in the data set to provide additional data for marine species and *Leporidae;* their isotopic data was obtained by O’Connell and colleagues (2019) at the nearby Portus site. The data distribution for the faunal remains is consistent with the expected locations in the food net and very few samples seem to be outliers. A bovine sample from Portus has a higher δ^15^N value than expected and the pigs from Ostia (Portus and Isola Sacra) and Colosseum area seem to suggest different foraging strategies due to their different enriched δ^15^N values. These could represent imported foodstuffs, and this could be consistent with the longstanding commercial connections between Rome and the nearby river and maritime ports of Portus and Ostia, as well as between Rome and other Mediterranean areas through the first centuries CE (O’Connel et al., 2019; Keay, 2013). Furthermore, the local baselines for Castel Malnome, Via Padre Semeria and Colosseum seem to roughly align with the ecological background determined for Portus and Isola Sacra for primary consumer herbivores. Accordingly, omnivores such as dogs from Rome lie one trophic level up and align with other Canidae from Isola Sacra and a bird from Portus. Unfortunately, no freshwater fish remains could be listed in the data set; while the diachronic marine fish values are accordingly located at less negative δ^13^C values (Figure 3).

**Figure 3:**
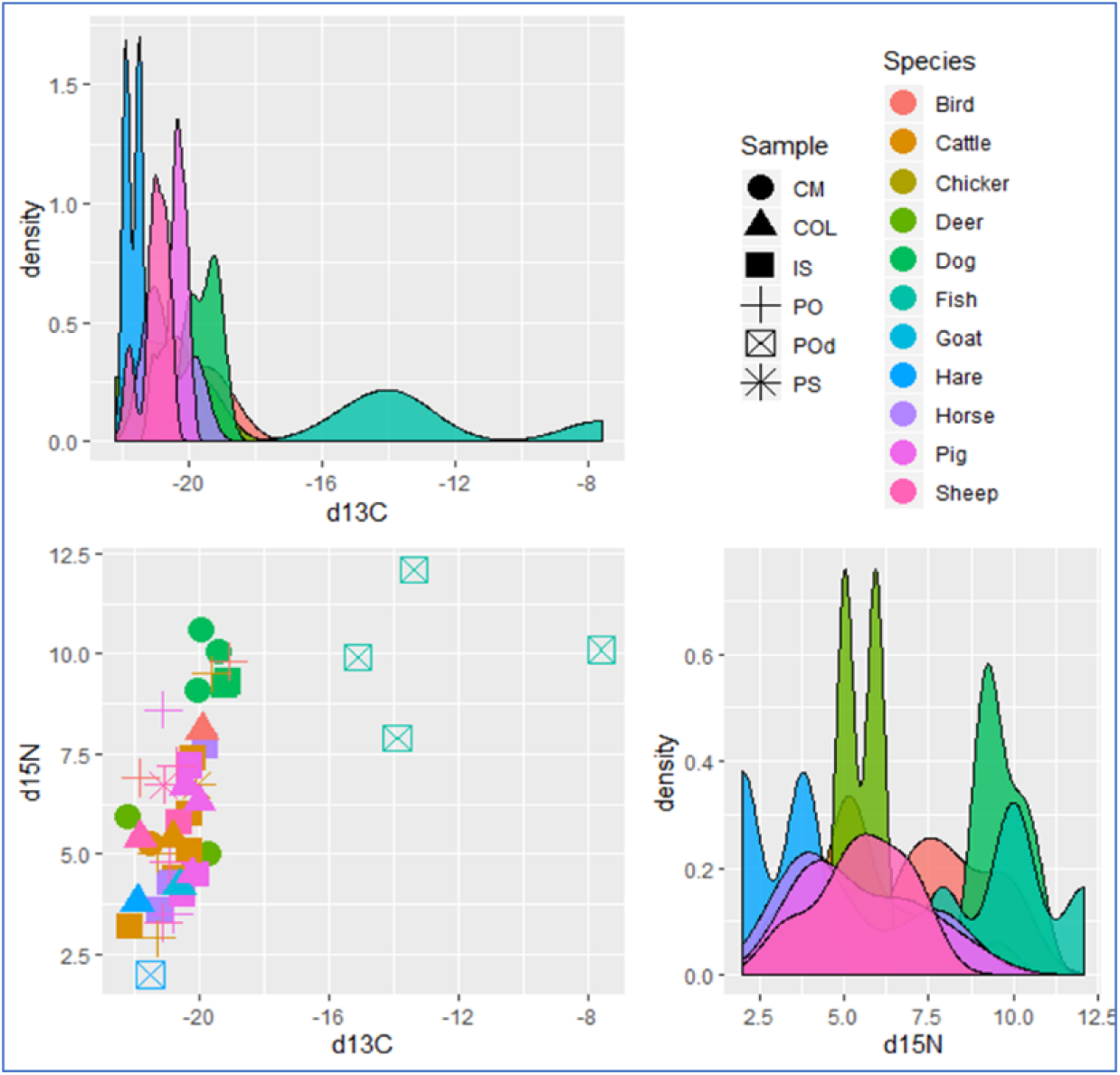
Bivariate distribution for faunal remains. CM: Castel Malnome, COL: Colosseum, IS: Isola Sacra, PO: Portus, POd: diachronic samples from Portus, PS: Via Padre Semeria.

The overall high trophic level of the humans (compared to the fauna) ensures the quality of the results and suggests that the livestock should be considered prey for the humans (Figure 4).

**Figure 4:**
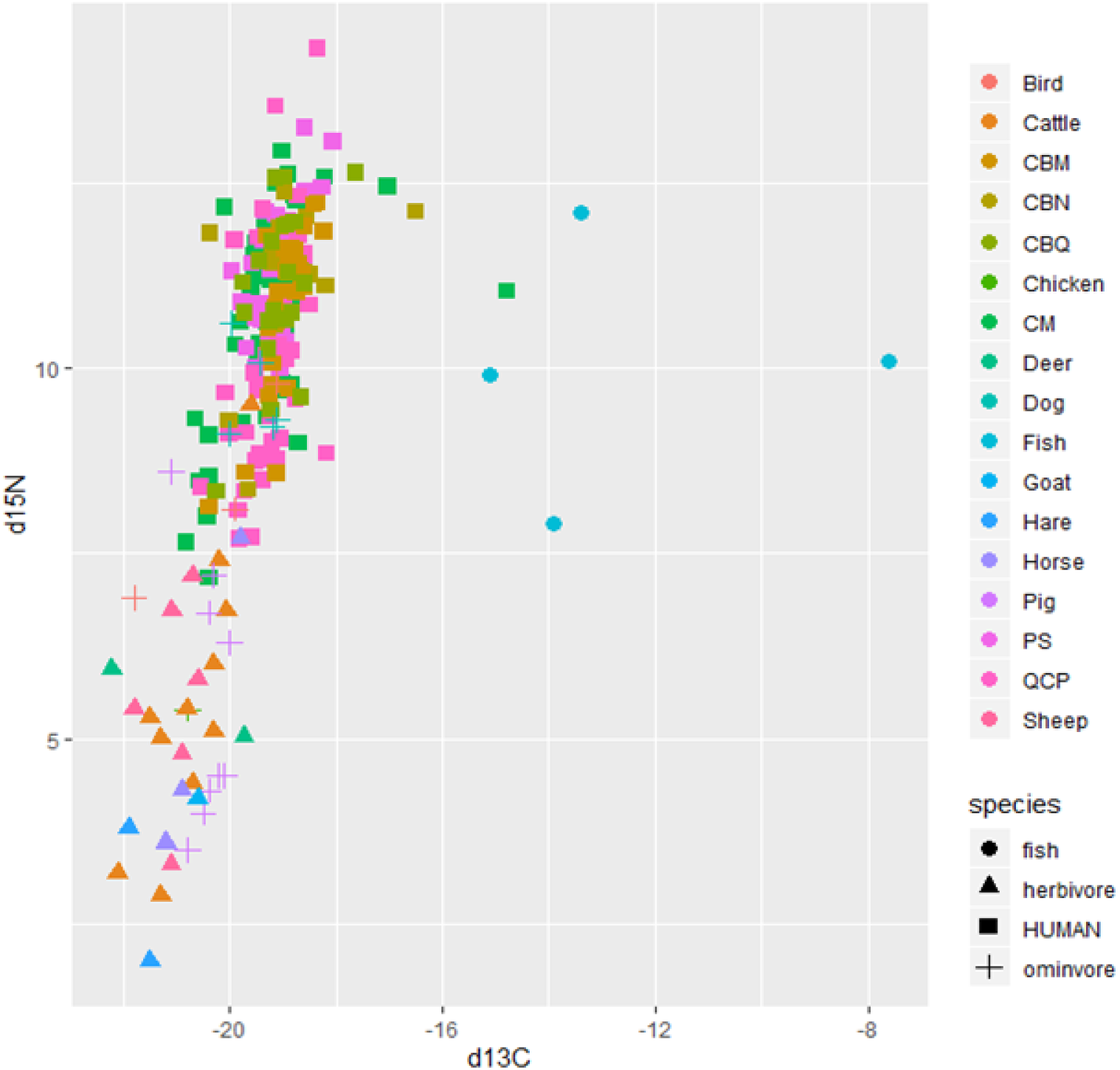
Plot for δ13C than δ15N values for humans and faunal remains. Color dot represent individuals: CBM, CBN, CBQ, CM, PS and QCP refer to humans from analyzed sites as reported in Table 1.

### DIET RECONSTRUCTION

The stable isotope analysis performed on the human remains recovered at the 6 funerary contexts in Imperial Rome (1^st^ – 3^rd^ CE) suggests people consumed a roughly heterogeneous diet based on C_3_ plant backbone resources. Several classical authors wrote about agricultural and horticultural practices in the Roman world: *Columella* (4^th^ BCE- 65^th^ CE) wrote *de Re Rustica*, which represents a valuable source of information on Imperial Roman agriculture, and Pliny the Elder is one of the best Roman sources on ancient plants. Although ancient literary sources on horticulture focused on the cultivation of olives and grapes for their significance in elite production (Lomas, 1993), these authors examined the production of cereal grains too, because they made up the bulk of most people’s diets as they were used to make bread and porridge (*puls*) (Brown, 2011).

At first glance, there is no evidence in the sample for exclusive C_4_ plant exploitation, supporting the notion that in Roman antiquity they were mainly consumed by animals rather than humans, even though the livestock data reported here does not suggest a foundational role for these plants.

Among these, millet represents a generic term for a large group of small-seeded grasses such as both *Setaria italica* and *Panicum miliaceum*. Millet is only mentioned a few times in ancient texts, and well-documented archaeological finds totally lack in archaeological surveys from Imperial Rome. Despite millet being not Romans’ first choice, it also was not totally discarded by the Romans, though it seems to have been more appreciated far from Rome. The presence of iconographic sources at estates in Pompeii suggests that millet may have been consumed by the wealthy landowners even though they did not totally appreciate it (Jashemski et al., 2002). Indeed, millet was found at several rustic Campanian and southern Italy estates (Boscoreale, Herolanum, and Matrice) (Spurr 1983, Murphy et al., 2013) and its role in cultual practices in northern Italy cannot be ruled out (Rottoli and Castiglioni 2011). Remarkably, millet was often noteworthy in relation to famines and food shortages (Spurr, 1983; Garnsey, 1999) due to its easy cultivation (Spurr, 1983): Columella reports that millet sustained the population of a lot of Italian provinces and the commercial value of millet in the Empire was set in the Edict of Diocletian. Even though millet, which unlike wheat is a non-glutinous grain, can make a heavy flat bread, this seed grass was preferentially used for animal fodder and birdseed rather than direct human consumption (Spurr, 1983). Nevertheless, millet was recommended for several medical uses, particularly for regulating the digestive system (Murphy et al., 2013), and its consumption seems to have been greater in several areas of the Roman empire. Although Killgrove and Tykot (2013) found a consistent use of C_4_ plants in Castellaccio Europarco for a small sample of buried people, bioarchaeological data about this cereal grain is scarce. Recent archaeobotanical evidence (O’Connel et al., 2019) shows a consistent amount of cereal grains, mainly free-threshing wheats, emmer, einkorn, and barley, at the Roman harbor of Portus, where no C_4_ plants were recovered. This direct evidence, though it could contain bias as a result of chance or for trade in the harbor, is consistent with the aforementioned Roman preference for C_3_ grains, along with pulses (lentils, peas, and broad beans were recovered) and fruits (a few grapes and elder berries were found in the flotation-sieved contexts at Portus).

The data distribution shows that there is no direct evidence of exclusive marine resource intake too. Although a few individuals, such as CM34, CM52, CBN1, and CBQ13, had positive values for δ^13^C, their moderate δ^15^N values do not clearly indicate marine fish consumption. The odd location could be due to a diet consisting of a combination of marine resources and mix of C_3_/C_4_ plant (or primary consumers who eat those plants) related to individual preferences and/or foodstuff availability. Nevertheless, their occasional consumption (along with freshwater resources) cannot be ruled out since up to 20% of the protein consumed could conceivably have come from marine ecosystems without any visible shift in collagen-derived values (Milner et al. 2004). For this reason, a mixed diet could easily be misidentified as an exclusively terrestrial diet (Jim et al. 2006). The seashore vicinity of Castel Malnome hints at the role marine resources could play in the diet, and a local creek could have provided supplemental freshwater food sources. Additionally, people buried at Casal Bertone could have accessed these resources through markets due to their proximity to the city walls.

The lack of evidence for exclusive marine fish (or shellfish) consumption should not be confused with infrequent consumption of fish through the Romans’ staple sauce, *garum*, which could be made with a variety of recipes. Much of the evidence about this ancient fish sauce comes from classical literary sources which postulate that its popularity derived primarily from social forces influencing individual tastes. The peculiar smell and taste made *garum* a popular food among wealthy people, although the general populace probably also used this condiment (Grainger, 2018). In addition, archaeological investigations are steadily uncovering evidence of the widespread production and sale of fish sauce throughout the Empire, suggesting that the wealthy were not the only ones with access to fish sauce (Grainger, 2018).

Recent archaeological findings in Portus turned up faunal remains that could represent a coeval dietary background (O’Connel et al., 2019). Sheep/pig-sized mammals made up the bulk of the findings, with the latter representing the most common species. Cattle, hares, and domestic fowl were also found, but very little venison. We are aware that these findings cannot fully represent the local foodstuffs for the communities buried in the analyzed necropolises, but by the same token, the evidence provided by the river harbor of Rome should not be undervalued.

Following the suggestions provided by Fontanals-Coll and colleagues (2016), we defined the average values in the dietary proxies for herbivores and omnivores from Rome and Portus/Ostia along with their variances, in order to draw the boxes where the prey could be set. These boxes are then shifted accounting for the predator-prey offsets, which have been estimated as +1‰ for δ^13^C and +4‰ for δ^15^N. These dietary markers increase with each trophic level and δ^15^N rises approximately +3/+5‰, with deviations depending on species and dietary composition, which suggest the use of the median value (Robbins et al., 2005; Fraser et al., 2013; Fontanals-Coll et al., 2016). The estimation of the consumers’ boxes indicates that most individuals fall inside the boxes built for herbivore consumers and omnivore consumers (Figure A.1) even though 48 individuals fall beyond the −18.67‰ threshold, indicating a clear C_3_-derived omnivore consumer (Table 7). This evidence pushes us to reconsider the fraction of people whose diet was based on mixed C_3_/C_4_ plants and/or marine resources. The data stratification for those 48 individuals by site, sex or age classes do not support any specific trend except for adult/child comparison for δ^15^N (δ^13^C: Kruskal-Wallis chi-squared = 8.44, df = 5, p-value = 0.13 for site; Kruskal-Wallis chi-squared = 0.03, df = 2, p-value = 0.98 for sex; Kruskal-Wallis chi-squared = 0.11, df = 1, p-value = 0.73 for age class; δ^15^N: Kruskal-Wallis chi-squared = 9.62, df = 5, p-value = 0.09 for site; Kruskal-Wallis chi-squared = 2.55, df = 2, p-value = 0.28 for sex; Kruskal-Wallis chi-squared = 4.40, df = 1, p-value = 0.04 for age class) suggesting a stratification between age classes in that sub-sample. People from Casal Bertone (both necropolis and mausoleum) seem to be overrepresented (33 out 48 people). The moderate δ^15^N values offset between humans and marine fish (mean values 11.48‰ for adults and 11.00‰ for children vs 10.00‰ for marine resources herein considered) might deter to exclusively consider this shift due to marine resources exploitation and this could be supported by notion that marine fish was considered expensive food in the Empire, suggesting that regular fish consumption may have been restricted to the upper strata of Roman society (Frayn, 1993). Additionally, the presence of several people buried in Casal Bertone (Musco et al., 2008) could advise to consider that they were a fairly wealthy group whose diet was varied and heterogeneous. This evidence is further supported by the topographical location of the cemetery, close to the city center and hence people buried in this area (and perhaps living and working at the same location, Catalano et al., 2015), could easily access to market system featuring the city of Rome, where several *horrea* (large warehouses and other storage facilities in Ancient Rome) were located (Vera, 2008; Burgers et al., 2015).

**Table 7:** Samples beyond C_3_ plant consumer threshold values

| Site | Sample | Sex | Age_class | $\delta^{13}\text{C}$ | $\delta^{15}\text{N}$ |
| --- | --- | --- | --- | --- | --- |
| **CBM** | CBM1 | M | Adult | -18.34 | 12.22 |
| **CBM** | CBM4 | M | Adult | -18.22 | 11.84 |
| **CBM** | CBM24 | M | Adult | -18.41 | 12.2 |
| **CBM** | CBM100 | F | Adult | -18.1 | 11.2 |
| **CBM** | CBM101 | M | Child | -18.1 | 10.7 |
| **CBM** | CBM102 | Ind | Child | -18.1 | 8.6 |
| **CBM** | CBM103 | Ind | Child | -18.1 | 10.3 |
| **CBM** | CBM104 | Ind | Child | -18.1 | 11.3 |
| **CBM** | CBM105 | F | Adult | -17.7 | 11 |
| **CBM** | CBM106 | M | Adult | -17.7 | 10.8 |
| **CBM** | CBM107 | F | Adult | -17.5 | 9.3 |
| **CBM** | CBM2 | F | Adult | -18.58 | 11.91 |
| **CBM** | CBM8 | Ind | Child | -18.61 | 11.08 |
| **CBM** | CBM9 | F | Adult | -18.63 | 11.26 |
| **CBN** | CBN1 | M | Adult | -16.49 | 12.1 |
| **CBN** | CBN2 | M | Adult | -18.2 | 11.1 |
| **CBN** | CBN100 | Ind | Child | -18.2 | 11 |
| **CBN** | CBN101 | M | Adult | -18.2 | 11.8 |
| **CBN** | CBN102 | M | Adult | -18.2 | 11.1 |
| **CBN** | CBN103 | M | Adult | -18.1 | 11.6 |
| **CBN** | CBN104 | Ind | Child | -18.1 | 9.8 |
| **CBN** | CBN105 | Ind | Child | -18.1 | 10.8 |
| **CBN** | CBN106 | F | Adult | -18.1 | 9.6 |
| **CBN** | CBN107 | M | Adult | -18.1 | 11.6 |
| **CBN** | CBN108 | Ind | Child | -18 | 10.8 |
| **CBN** | CBN109 | Ind | Child | -17.8 | 11 |
| **CBN** | CBN110 | Ind | Child | -17.5 | 13.2 |
| **CBN** | CBN111 | F | Adult | -17.4 | 10.2 |
| **CBN** | CBN112 | M | Child | -16.8 | 9.7 |
| **CBN** | CBN5 | M | Adult | -18.66 | 11.30 |
| **CBN** | CBN9 | Ind | Child | -18.49 | 11.25 |
| **CBN** | CBN11 | Ind | Child | -18.60 | 11.28 |
| **CBN** | CBN14 | M | Adult | -18.55 | 12.04 |
| **CBQ** | CBQ13 | F | Adult | -17.62 | 12.63 |
| **CBQ** | CBQ15 | F | Adult | -18.59 | 11.14 |
| **CBQ** | CBQ16 | M | Adult | -18.66 | 9.60 |
| **CM** | CM34 | M | Adult | -14.78 | 11.04 |
| **CM** | CM52 | M | Adult | -17.02 | 12.44 |
| **CM** | CM55 | M | Adult | -18.21 | 12.57 |
| **PS** | PS7 | F | Adult | -18.26 | 12.43 |
| **PS** | PS30 | M | Adult | -18.05 | 13.05 |
| **PS** | PS23 | M | Adult | -18.61 | 11.43 |
| **PS** | PS24 | M | Adult | -18.58 | 12.37 |
| **PS** | PS25 | F | Child | -18.58 | 13.23 |
| **QCP** | QCP25 | Ind | Child | -18.35 | 14.3 |
| **QCP** | QCP39 | M | Child | -18.17 | 8.84 |
| **QCP** | QCP2 | M | Adult | -18.58 | 11.53 |
| **QCP** | QCP16 | Ind | Child | -18.50 | 10.85 |

Unfortunately, no data pertaining to freshwater resources could be found in coeval and co-regional samples, so it could be tricky to model freshwater fish exploitation. Indeed, there is a paucity of archaeological evidence for the consumption of freshwater resources in the Empire because the archaeozoological record rarely includes lacustrine or riverine faunal remains and, when it does include them, they are in very small numbers and are difficult to obtain for analysis. Data pertaining to this kind of prey has been collected for two diachronic samples from pre-Roman Britain (Jay, 2008) and the late-Roman province of Pannonia (Hakenbeck et al., 2017). These data sets provide useful isotopic data concerning some freshwater resources. Despite the significant differences between the two samples (δ^13^C T-value: 3.10, p<0.01; δ^15^N T-value: 4.27, p<0.01), they consistently have low δ^13^C and high δ^15^N values. The data pertaining to Via Padre Semeria (δ^13^C median=19.24; mean=-19.13; var=0.2; δ^15^N median= 11.41; mean=11.38; var=0.76) and the necropolis location in the known hydrographic net of Almone river (Tallini et al., 2013) are consistent with a more than sporadic consumption of these foodstuffs (Figure 5), representing a supplement to a typical farming-derived diet. Indeed, the median value for people buried in Via Padre Semeria is shifted toward negative values for δ^13^C and positive values for δ^15^N (Figure A.2; Table 5), and even though the individual values do not fall within the boxes calculated for freshwater fish-consumers (Figure 5), a more than occasional use of these stocks cannot be denied.

**Figure 5:**
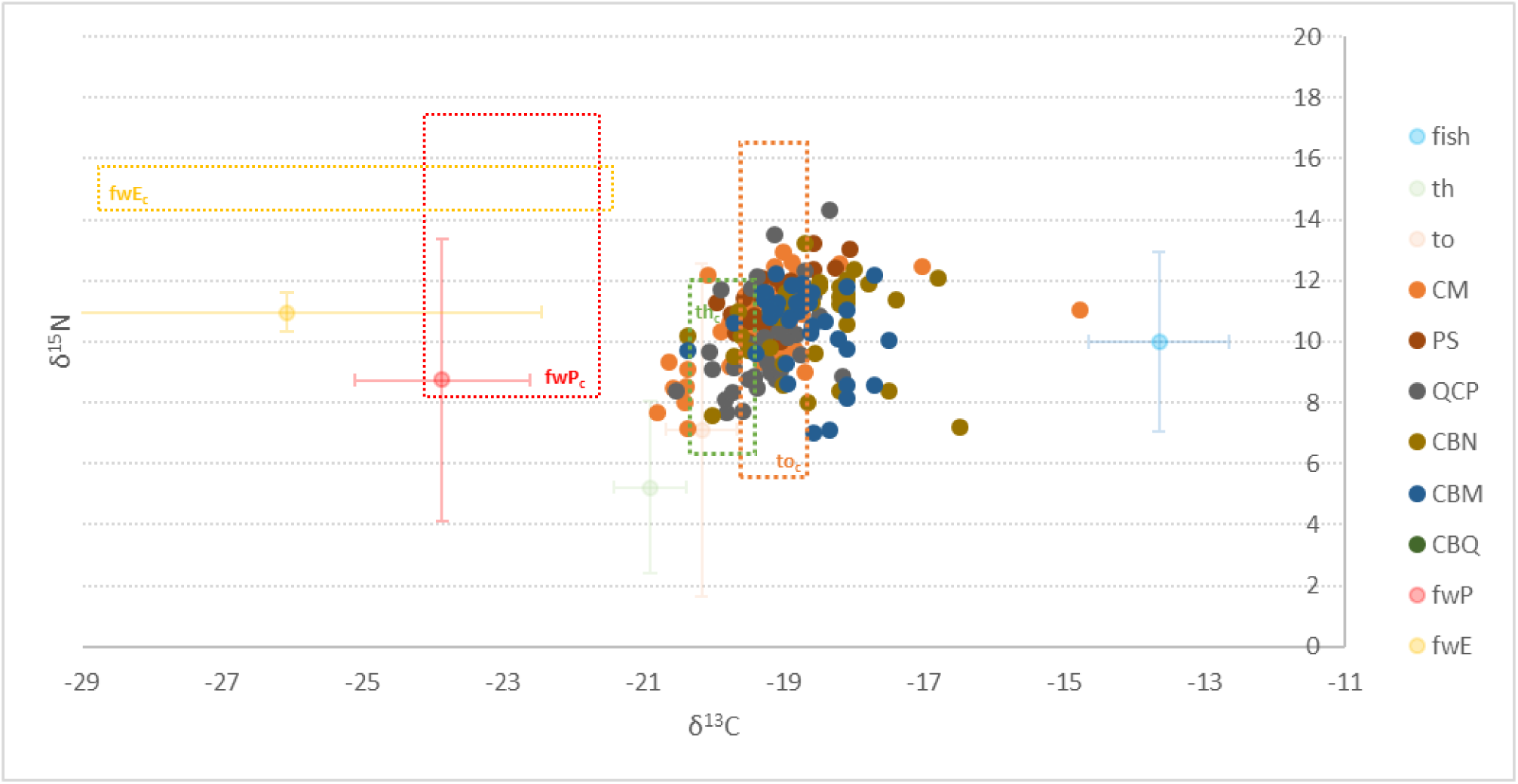
Linear model for identify prey-predator relationship. th: terrestrial herbivore; th_c_: terrestrial herbivore consumer; to: terrestrial omnivore; to_c_: terrestrial omnivore consumer; fwE_c_: freshwater fish form England consumers; fwP_c_: freshwater fish form Pannonia consumers; fish: marine fish. The dashed lines define consumers’ boxes.

Even though the median value determined for QCP is apart from other cemeteries (Figure A.2), suggesting a mostly farming-derived diet, the spread of the δ^13^C values collected for people buried in the cemetery appears to be meaningful (Table 5). This necropolis is related to a cultual site (Musco et al., 2001, Catalano, 2015) and mounting genomic evidence we are providing in an ancient DNA analysis for some individuals suggests people buried in it could be featured by genetic-related osteo-dystrophies (De Angelis, data not shown). Therefore, the necropolis could have served as a burial site for people suffering from skeletal diseases putatively coming from elsewhere, whose diets would be accordingly varied, as suggested by the wide δ^13^C range.

The median values calculated for CM and CBQ are quite similar (CBQ: δ^13^C =-19.13; δ^15^N=11.14; CM: δ^13^C =-19.16; δ^15^N=11.08) and this appears noteworthy due to the archaeological interpretations that these necropolises were tied to manufacturing activities (saltworks in CM and a tannery in CBQ). Meanwhile the necropolis and the mausoleum of Casal Bertone appear to be separate from CBQ, suggesting a certain degree of homogeneity that does not fit the archaeological data (Musco et al., 2008; Killgrove, 2013), which showed a clear difference in the social stratification of these samples: the isotopic results flatten the social mismatch, at least as concerns the dietary landscape.

### COMPARISONS

In order to fully explore the dietary scenario, we have attempted to understand the roles of several factors that could be significant in the onset of the differences among the necropolises.

All the data is normally distributed according to the Shapiro-Wilk Test (Table A.3), except for the Castel Malnome sample’s data (both δ^13^C and δ^15^N values) and the Casal Bertone mausoleum δ^15^N values, whose QQ-plots highlight their atypical distributions (Figure A.3, A.4 and A.5).

The data distribution for Castel Malnome suggests the presence of diet-based groups that could account for rough bimodal distributions of their δ^13^C and δ^15^N values, with the presence of some outliers: CM16 (male, 20-29 years old), CM20 (13-19 years, not available sex), CM21 (young female), CM23 (female, 30-39 years old), and CM33 (male, 40-49 years) feature significant low δ^13^C and δ^15^N values and those samples, along with CM66 (male, 30-39 years old), CM69 (male, 40-49 years old), and CM40 (male, 40-49 years old) seem to be clustered outside the normal distribution for δ^13^C, suggesting a plant-derived carbohydrate-rich diet. Conversely, CM34 (male, 20-29 years old), CM52 (male, 50 or more years old), and CM55 (male, 30-39 years old) are considered δ^13^C outliers that feature a more heterogenous diet. CM55, CM29 (male, 20-29 years old), and CM47 (male, 20-29 years old) are outliers for δ^15^N, suggesting a protein-rich diet.

The δ^15^N values in Casal Bertone mausoleum highlight a sub-stratification too. CBM 20 (child, 7- 12 years old), CBM 23 (child, 7-12 years old), CBM25 (teen, 13-19 years old), F10C (child, from Killgrove, 2010), F04B, and F11A (two adult females, from Killgrove, 2010) do not fit the normal distribution for low values, whereas CM1 (male, 30-39 years old), CM2 (female, 20-29 years old), and CM24 (male, 40-49 years old) are at the upper limit of the data distribution, falling outside the normal distribution.

The sample stratification was evaluated for the homoscedasticity through Levene’s test both for δ^13^C and δ^15^N and all the samples returned results that were not significant (δ^13^C Test statistic: 1.45, p=0.21; δ^15^N Test statistic: 1.48, p=0.20). The homogeneity of their variances allowed the Analysis of Variance (ANOVA) to detect differences among the groups.

This evaluation was performed taking into account site stratification, biological sex (male, female, or unknown due to skeletal immaturity), and age at death, with a dichotomic classification between adults and non-adults due to the variety of age classes reported, which was a result of different scoring methods used among the samples.

The ANOVA performed on δ^13^C values shows that the site represents a significant determinant for the onset of differences among the samples (F: 6.77, p= <0.01) while males, females, and children featured similar δ^13^C values (sex: F: 1.33, p= 0.27; age class: 0.45, p= 0.5).

The ANOVA performed on δ^15^N data shows that age class should represent an additional determinant for the onset of the differences among the samples (site: F: 4.00, p= <0.01; age class: F: 4.87, p= 0.03), while sex differences are negligible (sex: F: 0.90, p= 0.41).

The application of post-hoc tests is mandatory for the analysis of ANOVA results: they are designed for significant F-tests and additional exploration of the differences is needed to provide full information about the samples’ resulting differences from each other (Maxwell & Delaney, 2003).

The Tukey HSD test (Dubitzky et al., 2013) was performed to determine whether the relationship between two data sets is statistically significant. The obtained adjusted p-values account for multiple comparisons maintaining experiment-wise alpha at the specified level, set to 0.05 (Yuan and Maxwell, 2005). The application of *TukeyHSD* function in multcomp R package allows us to perform pairwise comparisons to obtain adjusted p-values that can dissect the real differences among samples. Accordingly, concerning δ^13^C, Castel Malnome seems split against Casal Bertone mausoleum, which is in contrast with Quarto Cappello del Prete. The same differences could be noted for Casal Bertone necropolis, which goes against the results for Via Padre Semeria (Table 8).

**Table 8:** Tukey HSD test results for δ^13^C according to Site. Asterisks indicate significant results.

| $\delta^{13}\text{C}/\text{Site}$ | | |
| --- | --- | --- |
|  | Test statistic | adjusted p |
| **CBN-CBM** | 0.13 | 0.95 |
| **CBQ-CBM** | -0.35 | 0.42 |
| **CM-CBM** | -0.48 | 0.01* |
| **PS-CBM** | -0.4 | 0.18 |
| **QCP-CBM** | -0.52 | 0.01* |
| **CBQ-CBN** | -0.49 | 0.10 |
| **CM-CBN** | -0.62 | 0.00* |
| **PS-CBN** | -0.53 | 0.02* |
| **QCP-CBN** | -0.65 | 0.00* |
| **CM-CBQ** | -0.13 | 70.97 |
| **PS-CBQ** | -0.04 | 1.00 |
| **QCP-CBQ** | -0.17 | 0.95 |
| **PS-CM** | 0.09 | 0.99 |
| **QCP-CM** | -0.04 | 1.00 |
| **QCP-PS** | -0.12 | 0.98 |

These results seem to be in accordance with qualitative results related to the medians (Figure A.2): CBM and CBN sit apart from the other necropolises, even though the range of the individual samples tends to homogenize the differences among the Casal Bertone areas and between Via Padre Semeria and CBQ and CBM.

In order to account for the differences in δ^15^N values, the same strategy was used, and the sole significant differences were found between Quarto Cappello del Prete against Castel Malnome and Via Padre Semeria, respectively (Table 9).

**Table 9.** Tukey HSD test results for δ^15^N according to Site. Asterisks indicate significant results.

| $\delta^{15}\text{N}/\text{Site}$ | | |
| --- | --- | --- |
|  | Test statistic | adjusted p |
| **CBN-CBM** | 0.14 | 1.00 |
| **CBQ-CBM** | 0.53 | 0.69 |
| **CM-CBM** | 0.30 | 0.85 |
| **PS-CBM** | 0.88 | 0.08 |
| **QCP-CBM** | -0.46 | 0.63 |
| **CBQ-CBN** | 0.39 | 0.89 |
| **CM-CBN** | 0.16 | 0.99 |
| **PS-CBN** | 0.74 | 0.20 |
| **QCP-CBN** | -0.60 | 0.33 |
| **CM-CBQ** | -0.23 | 0.98 |
| **PS-CBQ** | 0.35 | 0.94 |
| **QCP-CBQ** | -0.99 | 0.07 |
| **PS-CM** | 0.58 | 0.35 |
| **QCP-CM** | -0.76 | 0.04* |
| **QCP-PS** | -1.35 | 0.00* |

The post-hoc tests and the adjusted p-values determined for multiple testing allow us to exclude the previously identified differences noted between the age classes for δ^15^N (Test statistic: −0.20, p=0.27).

### COMPARISONS WITHIN ROME AREA

In an attempt to describe the food preferences of people buried in Rome, we collected the data for other funerary contexts such as Castellaccio Europarco (Killgrove and Tykot, 2013) and ANAS (Prowse, 2004) via IsoArch, and the data for people buried in nearby Isola Sacra (Prowse et al., 2001; Prowse, 2004, Crowe et al., 2010) and Portus Tenuta del Duca, O’Connel et al., 2019) are included for comparison.

The data evaluation by Levene’s test suggested there were dissimilarities in both δ^13^C and δ^15^N distributions’ variances (δ ^13^C: F value=3.97, p<0.01; δ ^15^N: F value=3.54, p=0.01) and thus we were forced to perform Kruskal-Wallis tests followed by post-hoc Dunn’s tests, the results of which were meaningful (Table A.4 and A.5).

The joint evaluation of the differences allows us to determine common patterns among necropolises, which could be grouped both on δ ^13^C and δ ^15^N axes according to the significant differences (the mean values of these can be seen in Figure 6 and 7, in the upper-right plots).

**Figure 6:**
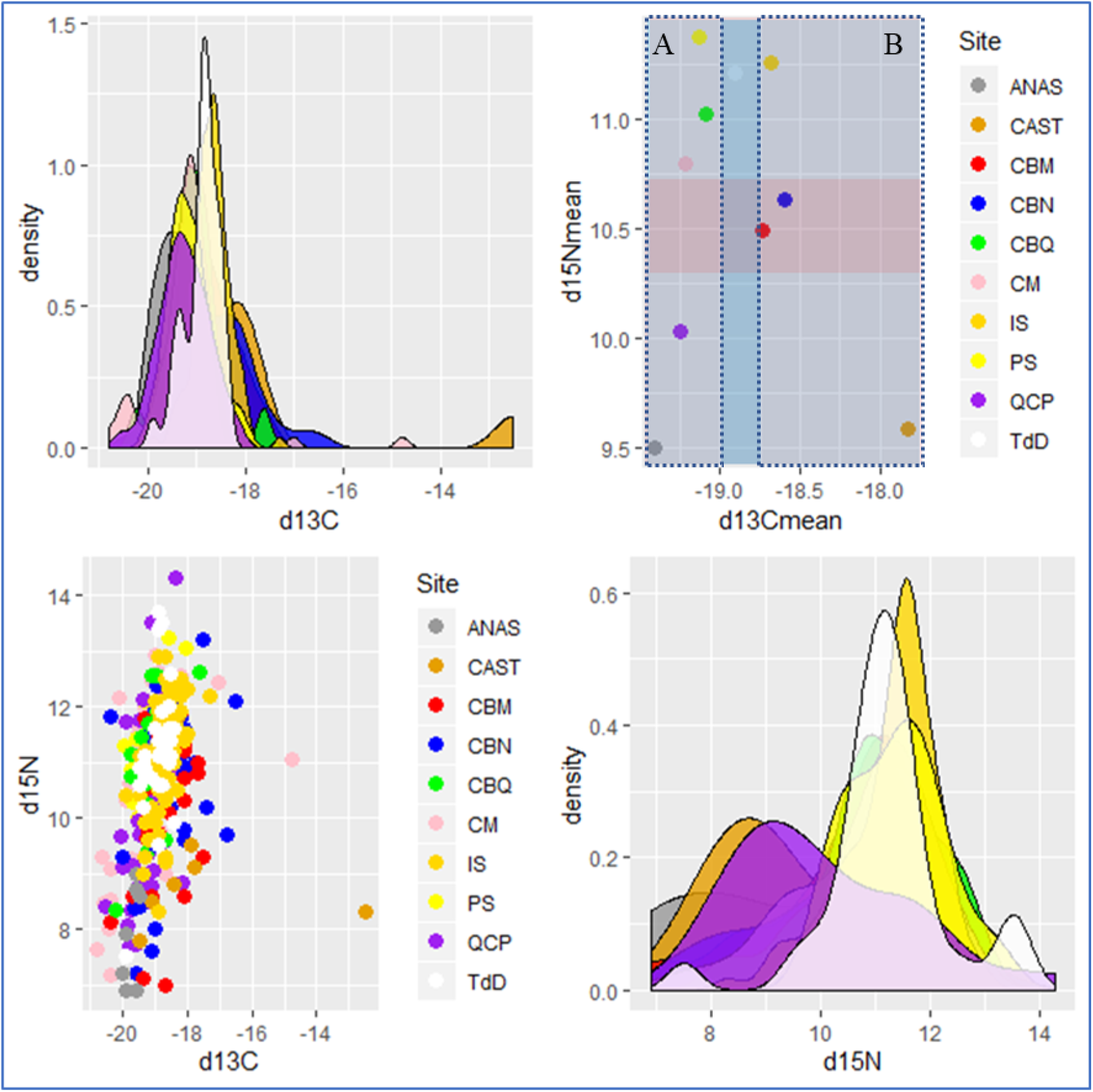
Data distribution, density plots and identification of A and B groups by joint Dunn’s Tests.

**Figure 7:**
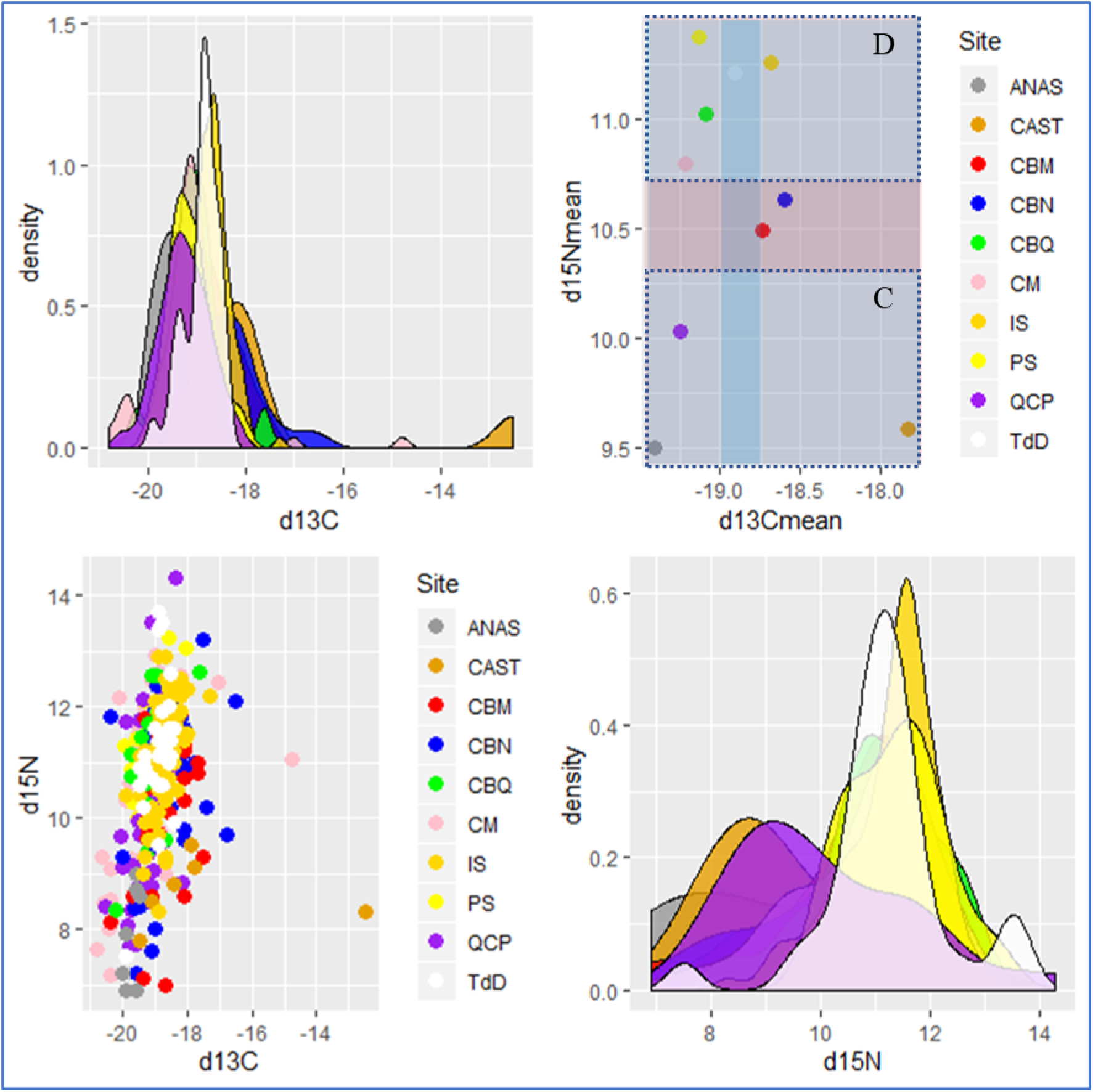
Data distribution, density plots and identification of C and D groups by joint Dunn’s Tests.

We can identify two segregated groups on the δ^13^C axis. One group (group A) comprises Via Padre Semeria, Casal Bertone Area Q, Castel Malnome, Quarto Cappello del Prete, and ANAS, which are significantly similar to each other and different from a second cluster (group B) made up of Isola Sacra, Casal Bertone Necropolis, and Castellaccio Europarco, with Tenuta del Duca (Portus) in the middle, sharing features from both group A and group B. This is only partially surprising if we consider the location of TdD close to the river harbor, where a lot of people from other areas and who ate different diets could have been buried.

Furthermore, δ^15^N values could be useful for determining two additional clusters. The first (group C) groups together ANAS, Castellaccio Europarco, and Quarto Cappello del Prete, while the second (group D) comprises Via Padre Semeria, Castel Malnome, Casal Bertone Area Q, and the sample from Portus. The low variance of Isola Sacra (var=0.77) could account for the sole exception to this clustering strategy based on significant similarities. Casal Bertone mausoleum and necropolis are located between these clusters, suggesting an intermediate condition.

This evidence could shed light on the overall dietary landscape for each site. The groups defined on the δ^13^C axis suggest that the individuals within them had different dietary habits. Group B represents people with a more heterogeneous diet, where C_4_ plant and marine resources consumption cannot be ruled out; this scenario is quite different from group A, which seems to have had a diet founded on C_3_ plants and freshwater fauna, that are consistent with the supposed diet reconstructions for Via Padre Semeria, Castel Malnome and Casal Bertone Area Q and Quarto Cappello del Prete.

The clusters on the δ^15^N axis should display meaningful differences in animal-derived protidic intake. Group C seems feature lower animal protein intake than group D, where terrestrial fauna along with lacustrine and riverine organisms likely provided needed dietary protein.

Nonetheless, Casal Bertone samples from the necropolis and mausoleum seem to feature intermediate characteristics and these are significantly different from area Q. The mausoleum and necropolis seem to be characterized by a more heterogeneous diet than area Q, with moderate protein intake, in which C_4_ plants as well as marine resource consumption cannot be totally ruled out. The archaeological differences (Musco et al., 2008) are not sustained by anthropological and isotopic evidence. Indeed, De Angelis and colleagues (2015) already recommended considering these contiguous areas as pertaining to a single population related to the *fullonica* that should be decoupled from area Q, where the demographic profile (and dietary habits too) suggests a separate community. Even though no certain archaeological data supports this hypothesis, it would be unusual for the funerary buildings of Area Q to be close to a productive area such as the tannery. In our opinion, we have to envisage the partial diachronic establishment of the tannery and the funerary buildings comprising Area Q.

The topographical location in the *Suburbium* (eastern suburbs, eastern suburbs close to city walls, south western suburbs, south western area close to Aurelian walls, Portus and Ostia) seems to indicate some differences (δ^13^C Kruskal-Wallis chi square=76.27, p<0.01; δ^15^N Kruskal-Wallis chi square=34.92, p<0.01) even though it is hard to identify a cline as well as some other common patterns. Nevertheless, the heterogeneous landscape highlighted in the Ostia and Portus samples are significantly different from eastern (Quarto Cappello del Prete and, close to the city walls, the whole sample of Casal Bertone) and southwestern (Castel Malnome, Castellaccio Europarco, ANAS, and, close to the city walls, Via Padre Semeria) necropolises (Table A.6 and A.7).

## CONCLUSIONS

The paper outlines the dietary landscape of Rome in the Imperial period. The evidence presented here provides a unique glimpse into the lives of the people who lived and died at Rome in established burial grounds, whose bio-cultural profiles have been described through detailed osteological, anthropological, and archaeological evaluations.

Despite the fact that the ancient Roman empire stretched across three continents, encompassing the Mediterranean region, the lives of tens of millions of people across Europe, the Near East, and North Africa (Antonio et al., 2019) were ruled by a centralized system based in Rome, where more than 1 million people crowded inside and outside the city walls. The necropolises located outside the walls are often related to individual communities, which were often made up of people of low social classes tied to productive sites or rustic *villae*.

The dietary landscape we provide is heterogeneous and this reflects the multifaceted reality of the capital of one of the most influential empires in Antiquity.

The complexity of Roman society remains difficult to disentangle even from a dietary point of view, but some elements can be illuminated. One of these is the pivotal role of C_3_ plants, which seem to have been the staple foodstuff of the lower class. However, C_4_ plants also seem to have been consumed, albeit they were not as widespread and were not always used for human consumption. The environment played a critical role also for Romans of lower social classes. Even though they were partially sustained by grain supplements from the central administration, the topographical location of the settlements (and perhaps of the necropolises where people were buried) determined the preferential consumption of food that people could obtain from their neighborhood, both farming and/or livestock breeding as well as by gathering (fishing and/or hunting), which provided the protidic intake needed to sustain active lifestyles defined by heavy labor. Nevertheless, the complexity of Roman society and trade that passed through Rome during the Imperial period accounted for the broader range of foodstuffs that people could access, making a portrait of the nutritional habits of Romans challenging, although the meticulous selection of burial grounds in this paper could lead to a less biased reconstruction. Indeed, exotic foods were only partially accessible to commoners, who mainly relied on local food resources, even though markets were accessible, especially to people living close to the city center. The proposed approach represents a powerful tool able to shed light on a crucial aspect of the biological characteristics of this ancient human population that pushes themselves beyond the biological feature: indeed, dietary patterns should be understood as one of the most long-lasting markers of the cultural identity of a population. The information provided herein represents a step forward in the understanding of the social organization of this ancient society, to be complemented by genomic and isotopic data related to migration, both in synchronic and diachronic perspectives (Killgrove and Montgomery, 2016; Antonio et al., 2019; De Angelis et al., in prep). The steady deepening of a combined archaeological and anthropological evaluation will allow us to stratify the Roman sample with respect to the bio- cultural factors that impacted the lives of a significant sample of the Roman populace.

## ACKNOWLEDGEMENTS

This work was supported by the Italian Ministry of Education, Universities and Research (MIUR) through PRIN 2015 (Diseases, health and lifestyles in Rome: from the Empire to the Early Middle Age, Grant ID: 2015PJ7H3K) allotted to CML, RSV and VG.

Authors would acknowledge Andy Bolduc and Martin Bennet for providing language help. Competing Interests statement: Authors declare that they have no significant competing financial, professional, or personal interests that might have influenced the performance or presentation of the work described in this manuscript.

## APPENDIX A

**Table A.1:** Data from Killgrove and Tykot (2013) related to Casal Bertone mausoleum and necropolis.

| Site | Sex | Age | Age_class | $\delta^{13}\text{C}$ | $\delta^{15}\text{N}$ |
| --- | --- | --- | --- | --- | --- |
| CBN | M | Adult | Adult | -19.6 | 7.2 |
| CBN | M | Adult | Adult | -19.5 | 8.4 |
| CBN | Ind | Non-adult | Child | -19.1 | 7.6 |
| CBN | M | Non-adult | Child | -19.0 | 9.5 |
| CBN | M | Adult | Adult | -19.0 | 8.0 |
| CBN | M | Adult | Adult | -18.6 | 11.3 |
| CBN | F | Adult | Adult | -18.5 | 10.2 |
| CBN | Ind | Non-adult | Child | -18.2 | 11.0 |
| CBN | M | Adult | Adult | -18.2 | 11.8 |
| CBN | M | Adult | Adult | -18.2 | 11.1 |
| CBN | M | Adult | Adult | -18.1 | 11.6 |
| CBN | Ind | Non-adult | Child | -18.1 | 9.8 |
| CBN | Ind | Non-adult | Child | -18.1 | 10.8 |
| CBN | F | Adult | Adult | -18.1 | 9.6 |
| CBN | M | Adult | Adult | -18.1 | 11.6 |
| CBN | Ind | Non-adult | Child | -18.0 | 10.8 |
| CBN | Ind | Non-adult | Child | -17.8 | 11.0 |
| CBN | Ind | Non-adult | Child | -17.5 | 13.2 |
| CBN | F | Adult | Adult | -17.4 | 10.2 |
| CBN | M | Non-adult | Child | -16.8 | 9.7 |
| CBM | F | Non-adult | Child | -19.4 | 7.1 |
| CBM | M | Adult | Adult | -18.7 | 7.0 |
| CBM | F | Adult | Adult | -18.6 | 11.0 |
| CBM | M | Adult | Adult | -18.6 | 10.1 |
| CBM | F | Adult | Adult | -18.1 | 11.2 |
| CBM | M | Non-adult | Child | -18.1 | 10.7 |
| **CBM** | Ind | Non-adult | Child | -18.1 | 8.6 |
| **CBM** | Ind | Non-adult | Child | -18.1 | 10.3 |
| **CBM** | Ind | Non-adult | Child | -18.1 | 11.3 |
| **CBM** | F | Adult | Adult | -17.7 | 11.0 |
| **CBM** | M | Adult | Adult | -17.7 | 10.8 |
| **CBM** | F | Adult | Adult | -17.5 | 9.3 |

**Table A.2:** isotopic values for faunal remains used in this paper. CM: Castel Malnome, PS: Via Padre Semeria, COL: Colosseum area, IS: Isola Sacra, PO: Portus. POd refers to diachronic samples from Portus (see text for details).

| **Sample** | **Species** | **$\delta^{13}\text{C}$** | **$\delta^{15}\text{N}$** | **Reference** |
| --- | --- | --- | --- | --- |
| **CM** | Dog | -19.4 | 10.1 | this paper |
| **CM** | Dog | -20.0 | 9.1 | this paper |
| **CM** | Dog | -20.0 | 10.6 | this paper |
| **CM** | Deer | -22.2 | 5.9 | this paper |
| **CM** | Deer | -19.7 | 5.0 | this paper |
| **CM** | Cattle | -21.5 | 5.3 | this paper |
| **PS** | Sheep | -21.1 | 6.7 | this paper |
| **PS** | Cattle | -20.1 | 6.7 | this paper |
| **COL** | Pig | -20.4 | 6.7 | this paper |
| **COL** | Goat | -20.6 | 4.2 | this paper |
| **COL** | Pig | -20.2 | 4.5 | this paper |
| **COL** | Chicken | -20.8 | 5.4 | this paper |
| **COL** | Bird | -19.9 | 8.1 | this paper |
| **COL** | Sheep | -21.8 | 5.4 | this paper |
| **COL** | Cattle | -20.8 | 5.4 | this paper |
| **COL** | Hare | -21.9 | 3.8 | this paper |
| **COL** | Pig | -20.0 | 6.3 | this paper |
| **IS** | Horse | -21.2 | 3.6 | Prowse et al., 2001 via IsoArch |
| **IS** | Cattle | -20.3 | 6 | Prowse et al., 2001 via IsoArch |
| **IS** | Horse | -19.8 | 7.7 | Prowse et al., 2001 via IsoArch |
| **IS** | Pig | -20.1 | 4.5 | Prowse et al., 2001 via IsoArch |
| **IS** | Pig | -20.5 | 4 | Prowse et al., 2001 via IsoArch |
| **IS** | Cattle | -22.1 | 3.2 | Prowse et al., 2001 via IsoArch |
| **IS** | Cattle | -20.3 | 5.1 | Prowse et al., 2001 via IsoArch |
| **IS** | Cattle | -20.7 | 4.4 | Prowse et al., 2001 via IsoArch |
| **IS** | Cattle | -20.2 | 7.4 | Prowse et al., 2001 via IsoArch |
| **IS** | Dog | -19.2 | 9.2 | Prowse et al., 2001 via IsoArch |
| **IS** | Pig | -20.3 | 7.2 | Prowse et al., 2001 via IsoArch |
| **IS** | Sheep | -20.6 | 5.8 | Prowse et al., 2001 via IsoArch |
| **IS** | Dog | -19.1 | 9.3 | Prowse et al., 2001 via IsoArch |
| **IS** | Horse | -20.9 | 4.3 | Prowse et al., 2001 via IsoArch |
| **PO** | Cattle | -21.3 | 5 | O'Connel et al., 2019 |
| **PO** | Cattle | -21.3 | 2.9 | O'Connel et al., 2019 |
| **PO** | Cattle | -21.3 | 5.0 | O'Connel et al., 2019 |
| **PO** | Cattle | -19.6 | 9.5 | O'Connel et al., 2019 |
| **PO** | Sheep | -20.9 | 4.8 | O'Connel et al., 2019 |
| **PO** | Sheep | -21.1 | 3.3 | O'Connel et al., 2019 |
| **PO** | Sheep | -20.7 | 7.2 | O'Connel et al., 2019 |
| **PO** | Pig | -21.1 | 8.6 | O'Connel et al., 2019 |
| **PO** | Pig | -20.4 | 4.3 | O'Connel et al., 2019 |
| **PO** | Pig | -20.8 | 3.5 | O'Connel et al., 2019 |
| **PO** | Bird | -19.1 | 9.8 | O'Connel et al., 2019 |
| **PO** | Bird | -21.8 | 6.9 | O'Connel et al., 2019 |
| **POd** | Hare | -21.5 | 2.0 | O'Connel et al., 2019 |
| **POd** | Fish | -13.9 | 7.9 | O'Connel et al., 2019 |
| **POd** | Fish | -15.1 | 9.9 | O'Connel et al., 2019 |
| **POd** | Fish | -13.4 | 12.1 | O'Connel et al., 2019 |
| **POd** | Fish | -7.6 | 10.1 | O'Connel et al., 2019 |

**Figure A.1:**
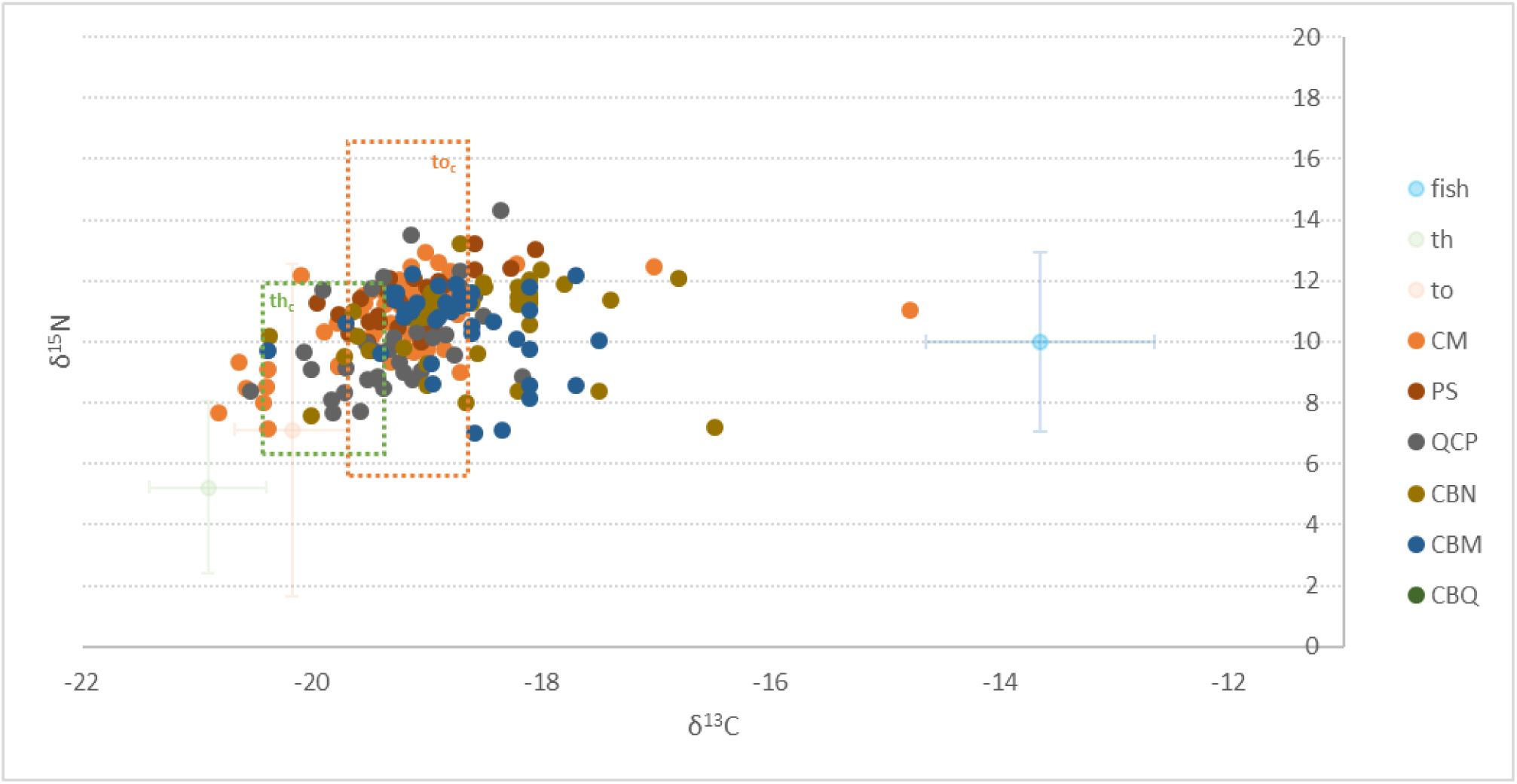
th: terrestrial herbivore; th_c_: terrestrial herbivore consumer; to: terrestrial omnivore; to_c_: terrestrial omnivore consumer. The dashed lines define consumers’ boxes obtained by the shifting of shaded lines representing the variances. Shaded dots represent the mean values for faunal resources.

**Figure A.2:**
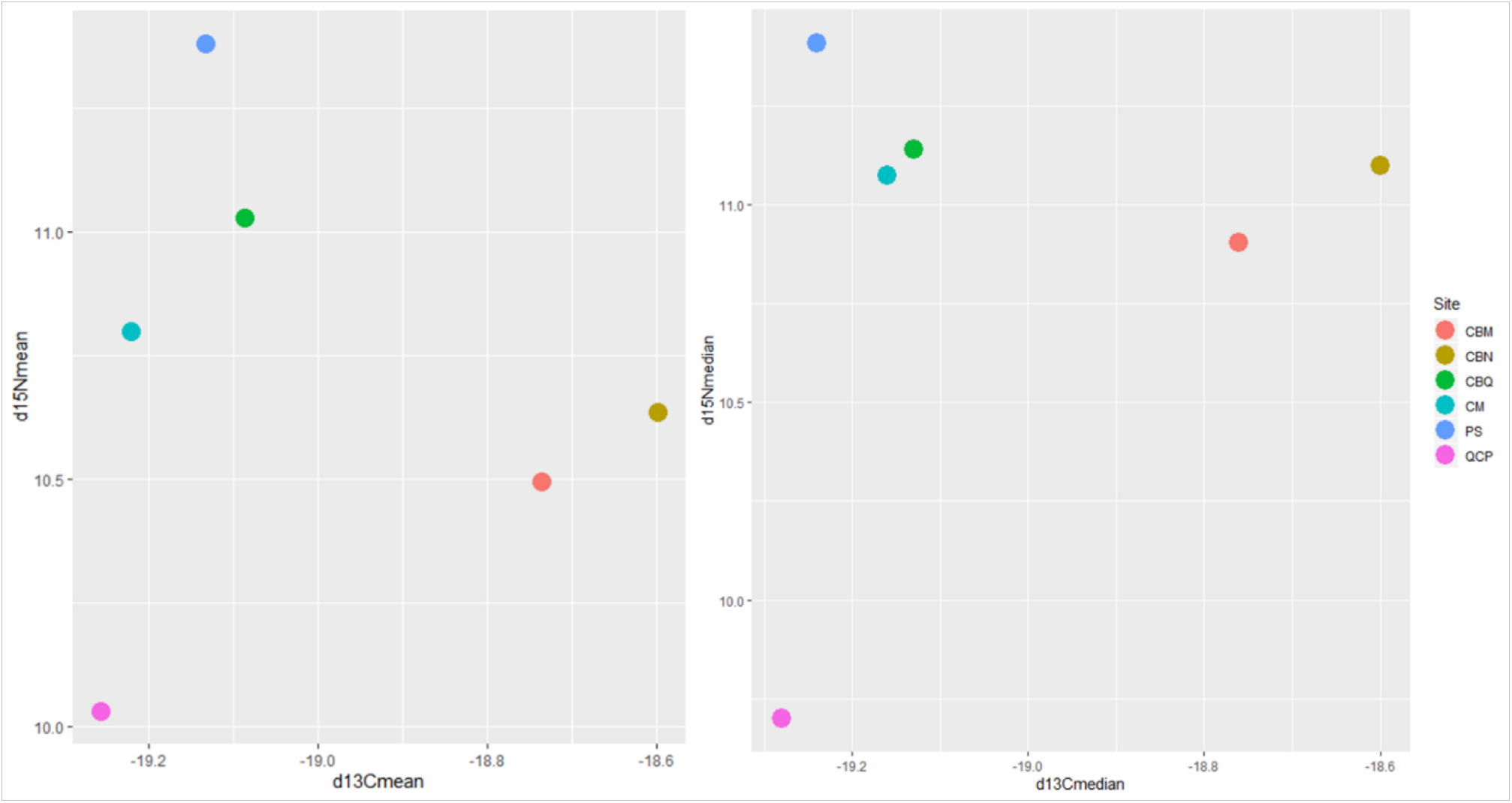
Mean (left) and Median (right) values distribution for each necropolis.

**Table A.3:** Shapiro-Wilk Test results.

| | $\delta^{13}\text{C}$ | | $\delta^{15}\text{N}$ | |
| --- | --- | --- | --- | --- |
|  | W | p-value | W | p-value |
| **CM** | 0.75 | <0.01 | 0.95 | <0.01 |
| **PS** | 0.97 | ns | 0.97 | ns |
| **QCP** | 0.99 | ns | 0.94 | ns |
| **CBN** | 0.98 | ns | 0.92 | ns |
| **CBM** | 0.97 | ns | 0.89 | <0.01 |
| **CBQ** | 0.93 | ns | 0.95 | ns |

**Figure A.3:**
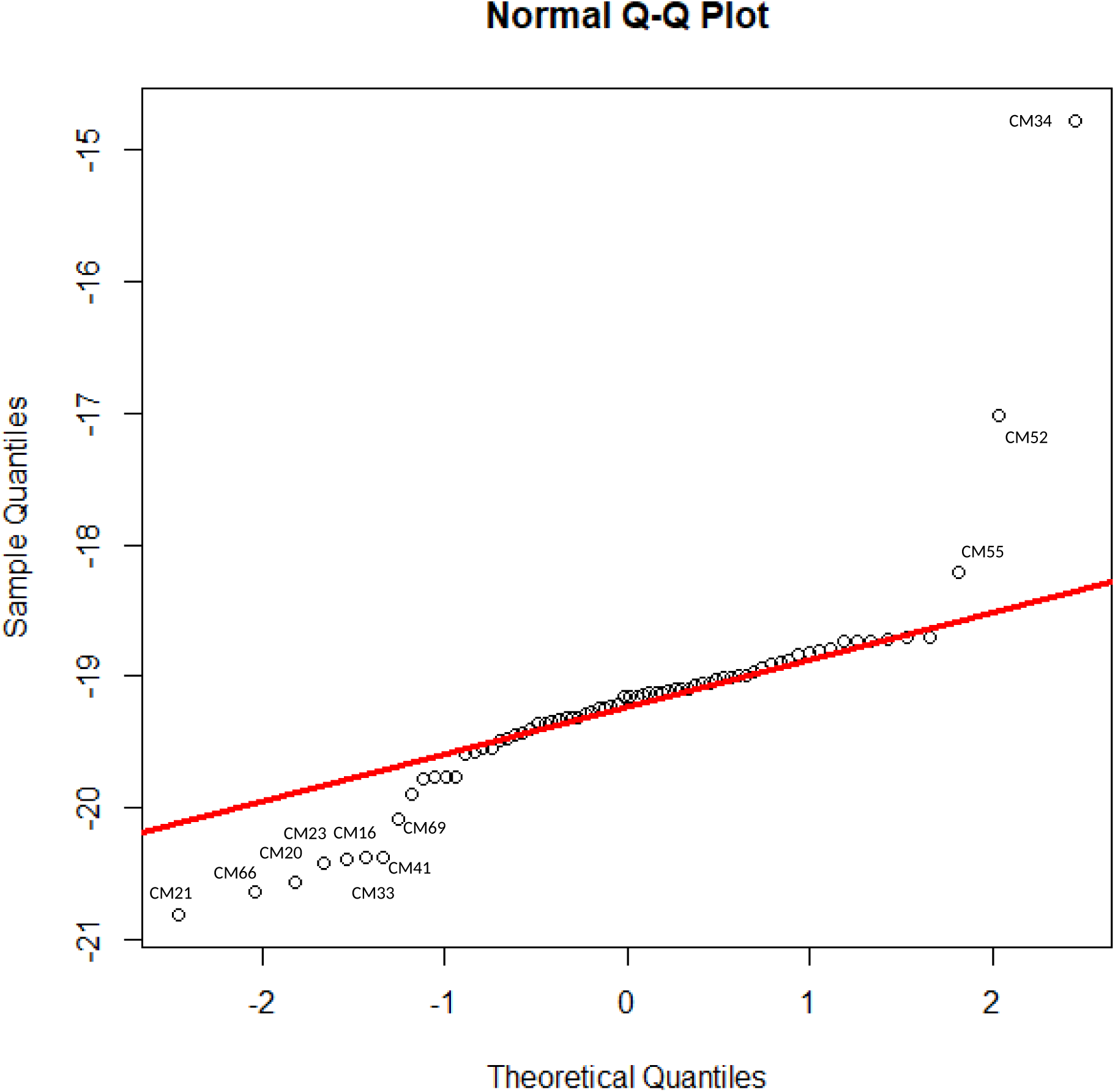
QQ plot defined for Castel Malnome δ^13^C values.

**Figure A.4:**
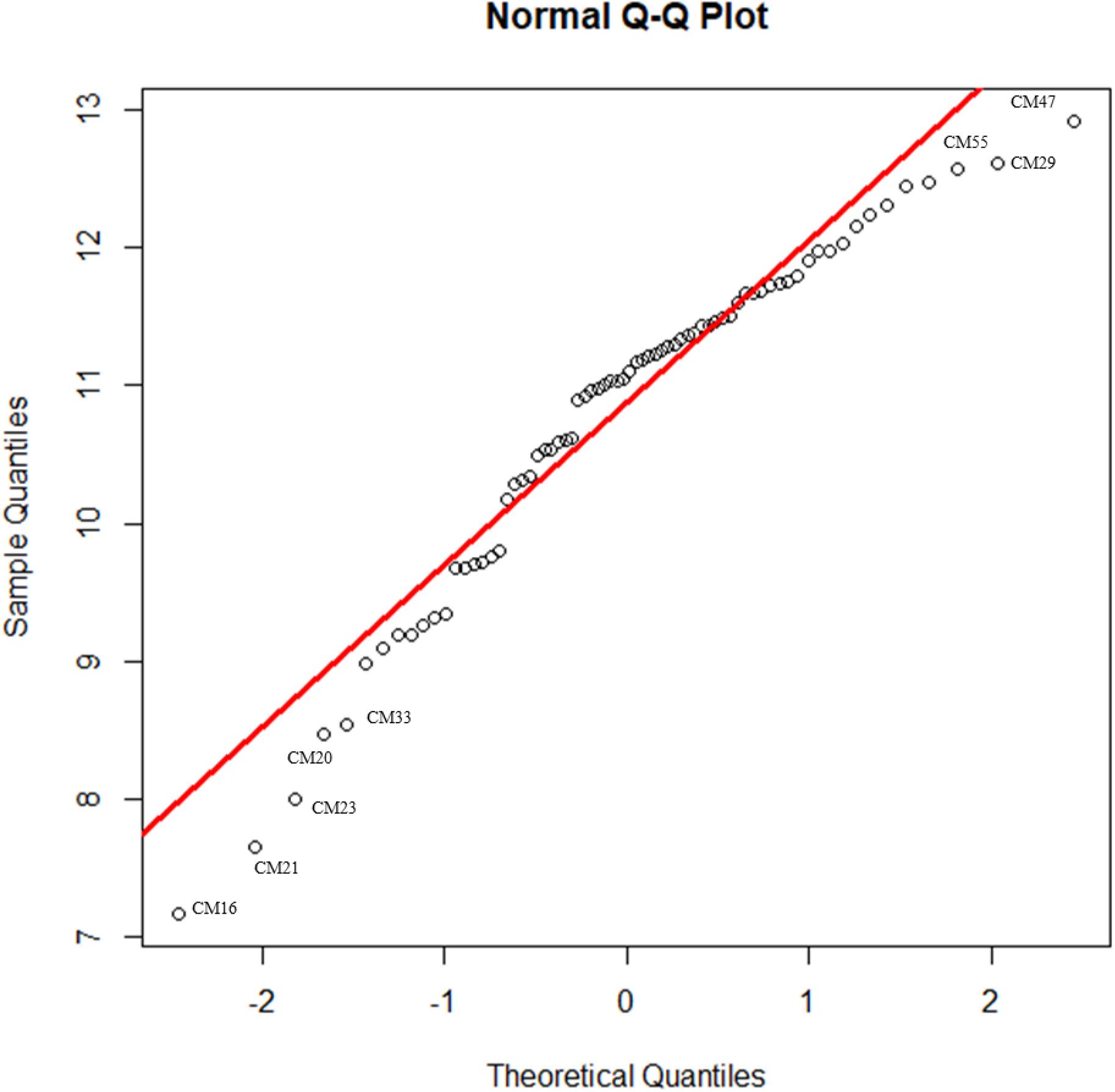
QQ plot defined for Castel Malnome δ^15^N values.

**Figure A.5:**
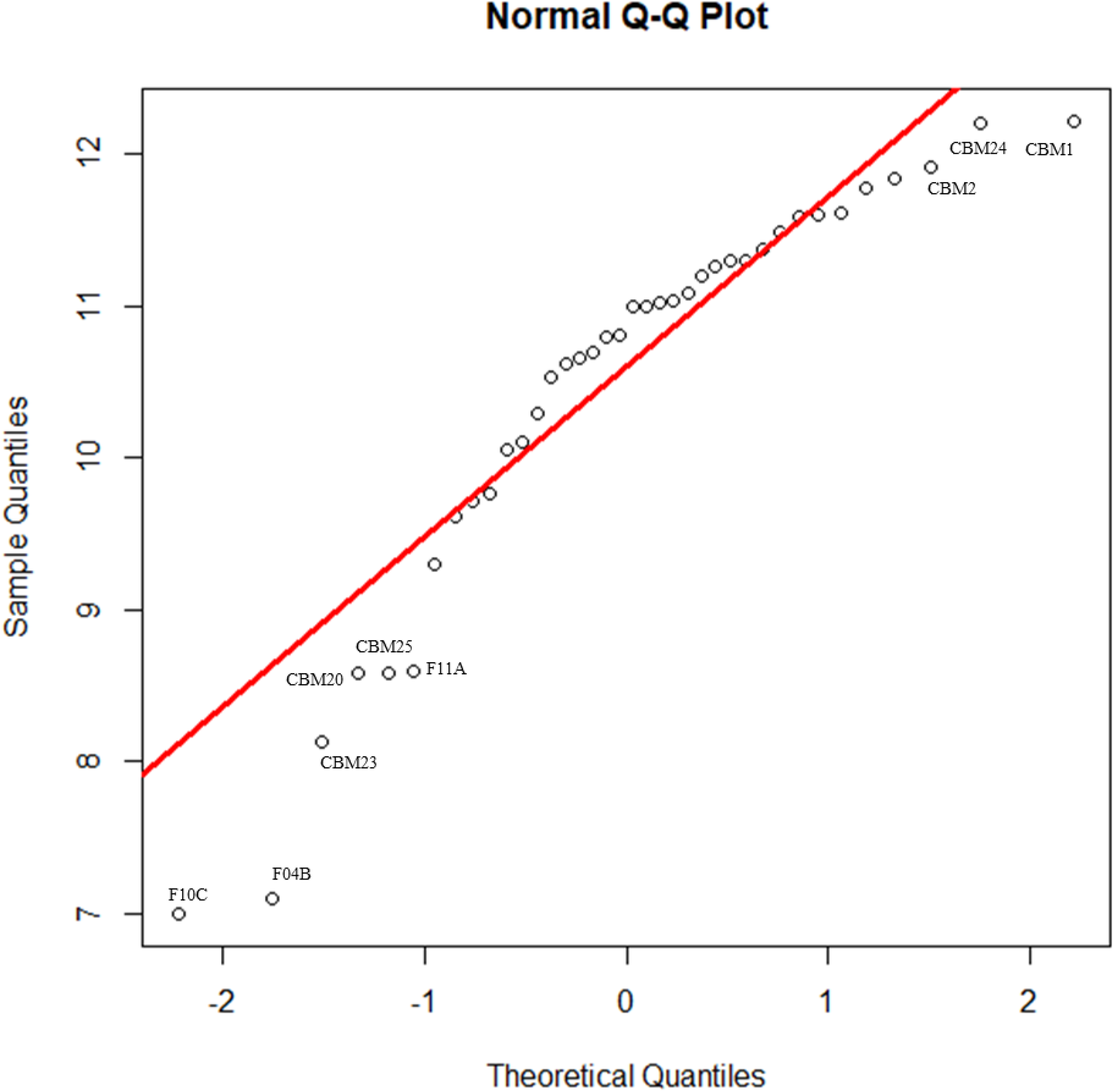
QQ plot defined for Casal Bertone mausoleum δ^15^N values.

**Table A.4:** Dunn’s test for δ ^13^C.

|  |  |  |  |  |  |  |  |  |  |
| --- | --- | --- | --- | --- | --- | --- | --- | --- | --- |
| Dunn's Test |  |  |  |  |  |  |  |  |  |
| **$\delta^{13}\text{C}$** | **ANAS** | **CAST** | **CBM** | **CBN** | **CBQ** | **CM** | **IS** | **PS** | **QCP** |
| **CAST** | -4.13 |  |  |  |  |  |  |  |  |
| **CBM** | -4.03 | 1.37 |  |  |  |  |  |  |  |
| **CBN** | -4.39 | 0.06 | -0.49 |  |  |  |  |  |  |
| **CBQ** | -1.72 | 2.86 | 2.33 | 2.73 |  |  |  |  |  |
| **CM** | -1.15 | 4.05 | 4.61 | 5.17 | 1.05 |  |  |  |  |
| **IS** | -5.15 | 0.83 | -1.12 | -0.54 | -3.46 | -7.27 |  |  |  |
| **PS** | -1.5 | 3.3 | 3.04 | 3.48 | 0.37 | -0.71 | 4.49 |  |  |
| **QCP** | -1.09 | 3.83 | 3.97 | 4.45 | 0.93 | -0.04 | 5.83 | 0.6 |  |
| **TdD** | -3.28 | 1.9 | 0.87 | 1.33 | -1.54 | -3.37 | 2.07 | -2.07 | -2.93 |

**Table A.5:** Dunn’s test for δ ^15^N.

| Dunn's Test |  |  |  |  |  |  |  |  |  |
| --- | --- | --- | --- | --- | --- | --- | --- | --- | --- |
| $\delta^{15}\text{N}$ | ANAS | CAST | CBM | CBN | CBQ | CM | IS | PS | QCP |
| CAST | -0.01 |  |  |  |  |  |  |  |  |
| CBM | -1.69 | -1.42 |  |  |  |  |  |  |  |
| CBN | -2.20 | -1.85 | -0.70 |  |  |  |  |  |  |
| CBQ | -2.52 | -2.19 | -1.28 | -0.71 |  |  |  |  |  |
| CM | -2.60 | -2.14 | -1.14 | -0.35 | 0.50 |  |  |  |  |
| IS | -3.99 | -3.27 | -3.20 | -2.37 | -1.02 | -2.46 |  |  |  |
| PS | -3.54 | -3.03 | -2.54 | -1.90 | -0.93 | -1.81 | -0.10 |  |  |
| QCP | -0.66 | -0.55 | 1.40 | 2.09 | 2.42 | 2.73 | 4.83 | 3.80 |  |
| TdD | -3.00 | -2.54 | -1.80 | -1.13 | -0.25 | -0.96 | 0.90 | 0.79 | -3.13 |

**Table A.6:** Dunn’s test for δ ^13^C according to topographical location. E: eastern suburbs, Ew: eastern suburbs close to city walls, SW: south western suburbs, SWw: south western area close to Aurelian walls, PO: Portus, OS: Ostia. Asterisks indicate significant results.

| $\delta^{13}\text{C}$ | E | Ew | OS | PO | SW |
| --- | --- | --- | --- | --- | --- |
| **Ew** | -4.29* |  |  |  |  |
| **OS** | -5.83* | -2.07* |  |  |  |
| **PO** | -2.93* | 0.6 | 2.07* |  |  |
| **SW** | -0.4 | 5.18* | 7.23* | 3.08* |  |
| **SWw** | -0.6 | 3.11* | 4.49* | 2.13* | -0.33 |

**Table A.7:** Dunn’s test for δ ^15^N according to topographical location. E: eastern suburbs, Ew: eastern suburbs close to city walls, SW: south western suburbs, SWw: south western area close to Aurelian walls, PO: Portus, OS: Ostia. Asterisks indicate significant results.

| $\delta^{15}\text{N}$ | E | Ew | OS | PO | SW |
| --- | --- | --- | --- | --- | --- |
| Ew | -2.37* |  |  |  |  |
| OS | -4.83* | -3.29* |  |  |  |
| PO | -3.12* | -1.45 | 0.9 |  |  |
| SW | -1.9 | 0.62 | 3.91* | 1.89 |  |
| SWw | -3.8* | -2.31* | -0.1 | -0.79 | -2.72* |

